# Enzyme-free biochemical production of seamlessly N-to-C cyclized peptides from natural or recombinant proteins

**DOI:** 10.1101/2025.07.31.667801

**Authors:** Ali Behboodian, Eugene Serebryany

## Abstract

Many naturally occurring or synthetic cyclic peptides are valuable as pharmaceuticals, but this stable and versatile class of molecules has not yet found applications beyond medicine. The main reason is the high cost of developing, producing, and altering these molecules via the gold-standard solid-phase synthesis methods. We focus on a class of cyclic peptides that have no disulfides, only canonical amino acids, and seamless peptide backbones. Known as orbitides or circular bacteriocins, such compounds are ribosomally synthesized and enzymatically cyclized by plants and bacteria. We report a simple method for producing them from naturally abundant proteins or from recombinantly expressed precursor polypeptides. The reaction proceeds under mild aqueous conditions, without the need for enzymes, and using only one chemical reagent, which is readily available. We demonstrate production of a 17-mer cyclic peptide from a wild-type human eye lens γ-crystallin and of a set of 10-residue cyclic peptides from recombinantly expressed polypeptide precursors. We investigate the effects of reaction conditions and sequence changes on reaction efficiency, identify the products by their complex mass spectrometry fragmentation patterns, and chromatographically separate linear and cyclic peptide forms. Our methodology opens the way to large-scale, cost-effective production of stable yet biodegradable, easily designable cyclic peptides for applications not only in medicine, but in areas like biotechnology, materials, agriculture, and pest control. It may also enable production of diverse cyclic peptide libraries from arbitrarily chosen natural protein sources.

## Introduction

Cyclic peptides (or peptide macrocycles) have attracted great interest and become widely used in drug discovery. They have many advantages over their linear counterparts: higher proteolytic stability, better cell permeability, and much lower conformational heterogeneity often leading to higher affinity and specificity (Ji et al., 2024). Owing to their constrained conformation and small number of epitopes, they are also less immunogenic than larger biologics, and perhaps even linear peptides (Driggers et al., 2008). Yet, they are large enough to target protein interfaces and other sites considered “undruggable” by conventional small-molecule pharmaceuticals (Naylor et al., 2017). Several classes of ribosomally synthesized, seamlessly head-to-tail cyclized cyclic peptides are known, e.g., orbitides and circular bacteriocins. Terminology is evolving and variable (Göransson et al., 2012; Ramalho et al., 2018; Simons et al., 2020; Tan & Zhou, 2006). Orbitides are characterized by seamlessly circular peptide backbone structure (i.e., cyclized by a backbone peptide bond), no side-chain crosslinks, and no non-canonical amino acids. This distinguishes them from non-ribosomally synthesized cyclic peptides, such as vancomycin.

Cyclic peptide binders for a target of interest can be discovered by high-throughput screening of designed libraries. The cyclic peptide libraries, in turn, may be generated in a variety of ways (Foster et al., 2015). Among the most popular are split-and-pool synthesis with DNA tags (Gartner et al., 2004; Usanov et al., 2018) and *in vitro* ribosomal synthesis and mRNA display (Goto & Suga, 2021; Huang et al., 2018). These approaches yield large and highly diverse cyclic peptide libraries, but tagging the peptides with much larger nucleic acid molecules (RNA, DNA, or both) to enable efficient identification of hits at the very small scale of these experiments. Nucleic acid tags, in turn, further limit the scope of biophysical experimentation: for example, they interfere with assays for membrane permeability (Dougherty et al., 2019; Faris et al., 2024); moreover, any study of pharmacokinetics *in vivo* using tagged peptide libraries is likely to produce artifactual results due to the tags. Smaller libraries of tag-free constructs can be screened by mass spectrometry (Bruce et al., 2024; Lee et al., 2025). The alternative method is *in cellulo* generation and screening (Osher & Tavassoli, 2017; Tavassoli & Benkovic, 2007), which yields smaller libraries of only natural amino acid containing peptides and requires screening inside the bacterial cell. Lastly, owing to advances in physics-based (Hosseinzadeh et al., 2021) and deep learning-based computational modeling of protein-peptide interactions, *in silico* screening of large cyclic peptide libraries is becoming increasingly possible and successful (Daumiller et al., 2025; Rettie et al., 2025).

However, a major bottleneck for all these approaches is downstream of high-throughput screening or design: high production cost of the cyclic peptides at large scale. Designed cyclic peptides are currently produced by solid-phase synthesis, which typically costs hundreds of dollars per gram even for simple linear peptides (ABI, 2025). Further adding to the cost, the cyclization yields are often below 20% even for short peptides (Rosenbaum & Waldmann, 2001), although native chemical ligation methods can yield as much as 50% (Wierzbicka et al., 2021). High cost of cyclic peptide manufacturing is often acceptable for high-value pharmaceuticals, but it makes most applications beyond human therapeutics economically non-viable. For example, cyclic peptides in general and orbitides in particular could be safe-yet-stable biological pesticides that could be sprayed on fields, long-lasting yet fully biodegradable hormone analogues for animals, or readily tunable building blocks for biomaterials such as nanotubes (Abdalla & McGaw, 2018). In contrast to the hundreds of dollars per gram cost of solid-phase synthesis, recombinant proteins produced by industrial-scale fermentation cost only cents per gram (Cardoso et al., 2020). It is therefore logical to explore whether cyclic peptides can be produced from proteins derived from fermentation or from naturally abundant sources.

Considerable bioengineering efforts have focused on converting proteases such as subtilisin into peptide cyclases (Chang et al., 1994; Schmidt et al., 2017). A more bioinspired approach has focused on biotechnological applications of enzymes such as butelase I, naturally present in orbitide-producing plants (James et al., 2019; Nguyen et al., 2014). Yet another alternative is biochemical production via split-intein ligation(Peña et al., 2022). However, these methods rely on expensive and chemically labile enzymes, and/or they require engineering of the recombinantly produced precursor with unique sequences to enable cyclization.

Here we report a simple method for enzyme-free biochemical production of seamlessly N-to-C cyclized peptides from recombinantly generated designed linear polypeptide precursors or even from naturally occurring proteins not evolved for this purpose. We achieve cyclization by modifying the methodology initially developed for Cys-targeted protein cleavage (Gary R Jacobson et al., 1973; Serebryany et al., 2023; Wu & Watson, 1997) to achieve peptide cyclization instead. Our methodology entails only two chemical steps: (1) cyanylation of a Cys residue between the N-terminal peptide and the rest of protein molecule, and (2) intramolecular aminolysis via nucleophilic attack of the free amine group of the peptide N-terminus on the peptide bond immediately upstream of the cyanylated Cys residue, which serves as the leaving group (**Figure 1**). Despite its simplicity, the potential of this type of chemistry has been largely overlooked. To our knowledge, it has been used successfully to fuse a protein’s termini together (Takahashi et al., 2007) but never to excise orbitide-like (or any other) cyclic peptides from linear protein precursors. Producing protein precursors in bacteria and obviating the need for enzymes in the cyclization process maximizes robustness of our method and minimizes cyclic peptide production costs.

**Figure 1.**
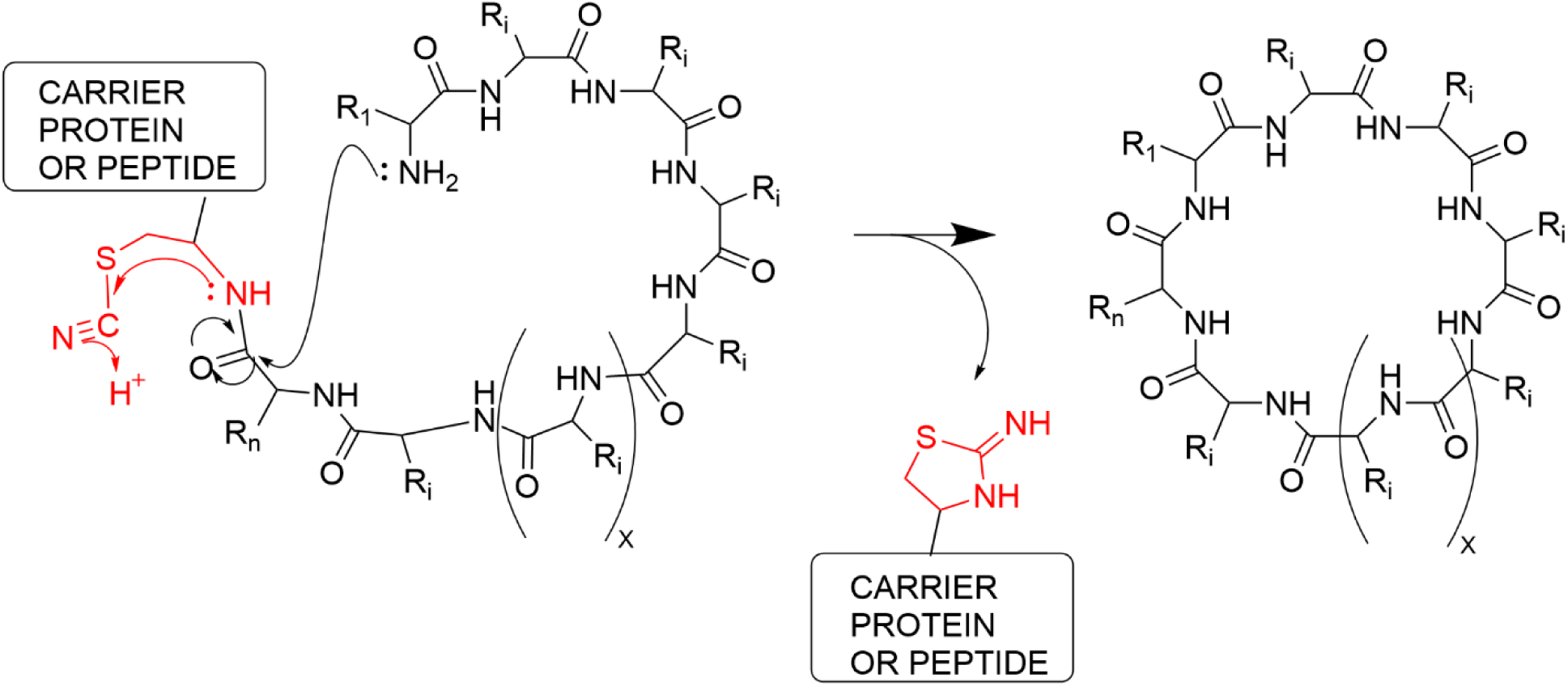
Schematic representation of the peptide cyclization reaction. In this approach, the Nitrogen of N-terminus amino acid attacks the peptide bond immediately upstream of the cyanylated Cysteine residue, with the carrier polypeptide becoming the leaving group and the N-terminal peptide upstream of the Cys residue becoming seamlessly N-to-C cyclized.

We demonstrate the applicability of this facile, enzyme-free approach to generating cyclic peptides either from naturally occurring proteins or from recombinant synthetic constructs. We demonstrate production of a 17-mer cyclic peptide from the N-terminus of wild-type human lens γD-crystallin (HγD), with the sequence c(GKITLYEDRGFQGRHYE). We also demonstrate production of a set of 10-mer cyclic peptides, sequence c(GKFISREFHX), where X = A, R, or E, from a recombinant fusion construct, in which the desired sequence is fused to the N-terminus of a small protein domain acting as a carrier. The resulting head-to-tail cyclzed peptide macrocycles are identified by mass spectrometry, which confirms cyclization by revealing diverse fragmentation patterns arising from the same cyclized precursor sequence. Finally, we explore the effects of varying the residues upstream and downstream of the cyclization site on the efficiency of initial cyanylation and final cyclization, with implications for the generalizability and yield optimization of this method.

## Results

### Cyclic peptide production from wild-type HγD crystallin

To validate the possibility of this approach on a natural protein, we took advantage of wild-type Human γD-crystallin (HγD), since it has a cysteine residue at position 18 in the mature form (**Figure 2a**). We unfolded this protein, at 1.9 mg/mL, overnight at 37 °C in 4M guanidinium chloride (GdnHCl) pH7 and 1 mM Tris(2-carboxyethyl)phosphine (TCEP) (final concentrations) to ensure a fully unfolded structure and fully reduced Cys residues. We desalted the unfolded protein into 4M GdnHCl pH 3 without TCEP. Cyanylation was done by adding 1-Cyano-4-dimethylaminopyridinium (CDAP) (2mM final concentration) and incubating at room temperature for 3.5 hours. After the cyanylation step, the reaction was divided and desalted into three buffers (A:1M Phosphate buffer, pH 7; B: 1M Phosphate buffer, pH 8; C: 1M Carbonate buffer, pH 9). For each buffer, the aminolysis reaction was done by incubating at 4 °C vs. room temperature overnight. The samples were quenched by adding 0.1% formic acid and were analyzed by mass spectrometry. According to the mass spectrometry analysis, the best cyclization yield was observed when aminolysis step was done at pH 9, 4 °C overnight, which yielded linear and circular products in approximately 1:1 ratio based on total ion current (**Figure 2b**). The molecular mass of the cyclic product differs from that of the linear product by the mass of one water molecule (**Figure 2c**). To make sure the product was cyclic, rather than a dehydrated linear peptide, we carried out LC/MS/MS (**Figure 2d**). Unlike the linear peptide, which fragments only from the ends, the ring of cyclic peptide can open at multiple positions, some of which may be preferred due to secondary structure or sequence features. Indeed, inspection of the complex MS/MS pattern of the cyclic product readily revealed two major fragmentation patterns (**Figure 2d**). Notably, residues 1-18 form most of a β-hairpin in the native protein structure (Basak et al., 2003), and simulations have shown this hairpin to be exceptionally stable even in the absence of cyclization (Serebryany et al., 2016). Accordingly, in the MS/MS experiments, the peptide appears to open preferentially on either side of the newly formed peptide bond, which probably strains the hairpin structure.

**Figure 2:**
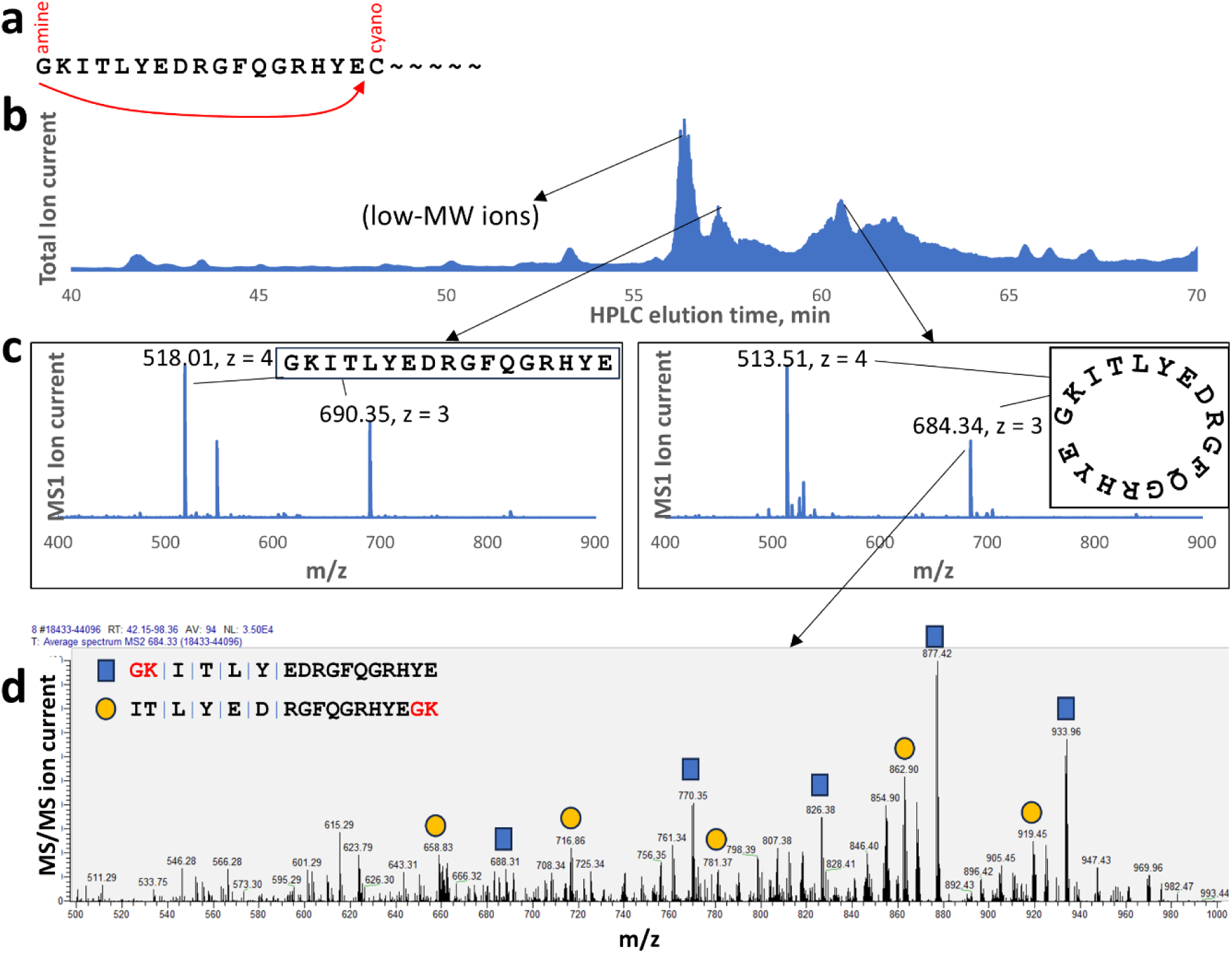
Production of linear and circular peptides from a natural protein via cyanylation-aminolysis chemistry. (a) Schematic of cyanylation-induced cyclization of the N-terminus of human γD-crystallin to produce (b) a mixture of peptides that include (c) the cyclic version of peptide 1-17 (*right*) and a competing linear hydrolysis product (*left*). MS/MS fragmentation of the cyclic product (d) yields a complex pattern indicative of quasi-random ring-opening; markers show two of the most prominent opening patterns.

### Design of a Cys- and Lys-free carrier domain for peptide-protein fusions

Encouraged by the observation of cyclic peptide production from wild-type HγD crystallin, we engineered its native C-terminal domain to attach a short peptide plus a cysteine residue to its N-terminus (**Fig. 3**). This crystallin domain is a particularly appropriate carrier for the short peptide due to its high stability and absence of Lys residues in its sequence (Mills-Henry et al., 2019). We also genetically removed both its native Cys residues, via the C108A/C110S substitution and appended a His_6_-tag to the C-terminus of the construct to facilitate affinity purification. The peptide-protein fusion construct was produced via conventional recombinant expression in *E. coli*. The short peptide’s sequence was chosen arbitrarily. We chose the sequence of a previously reported α-crystallin-derived mini-chaperone peptide FISREFHR (Raju et al., 2016), adding GK to its N-terminus to facilitate bacterial expression. Thus, the overall amino acid sequence of construct 1 was the following, with the sequence of the peptide underlined:

<u>GKFISREFHR</u>**C**GSRIRLYEREDYRGQMIEFTEDASSLQDRFRFNEIHSLNVLEGSWVLYEL SNYRGRQYLLMPGDYRRYQDWGATNARVGSLRRVIDFSHHHHHH

**Fig 3.**
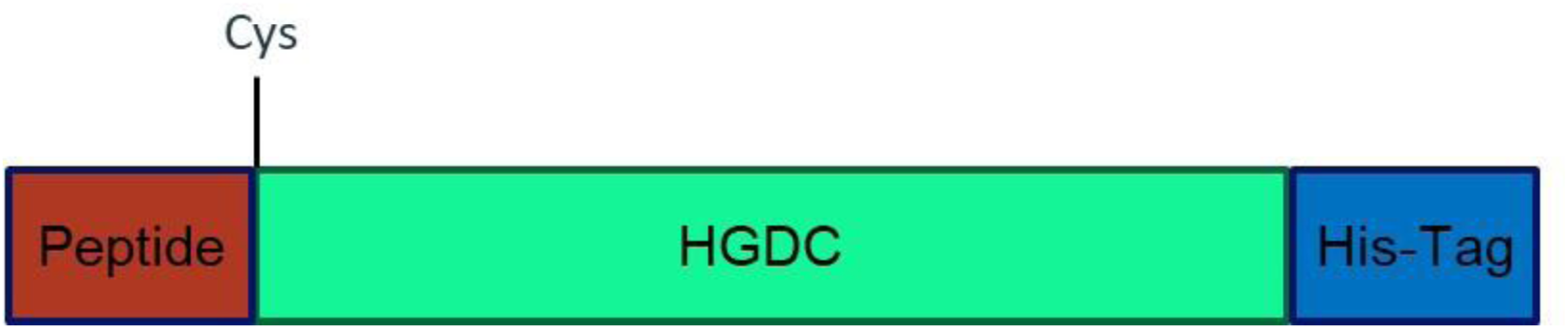
Design of the recombinant peptide-protein fusion constructs.

### Biochemical production of a cyclized mini-chaperone peptide

The purified construct containing the mini-chaperone peptide at the N-terminus was subjected to cyanylation and aminolysis similarly to the WT HγD crystallin above. However, since the recombinantly added peptide sequence was not part of the protein core, we found that protein denaturation was optional in this case. The pH was decreased by adding acetate buffer, and CDAP was used as the cyanylating agent, and the cyanylation-aminolysis reaction was performed as described in **Methods**. The mixture then was analyzed by SDS-PAGE to confirm and quantify overall reaction yield, and by MS to confirm production of the new cyclic peptide and determine the ratio of linear to cyclic products. We systematically varied several reaction parameters to optimize the efficiency of both reaction steps: cyanylation and aminolysis.

### Optimization of the cyanylation and aminolysis reactions

The overall efficiency of the cyanylation-aminolysis reaction can be readily monitored and quantified by SDS-PAGE, comparing the intensity of the cleaved vs. uncleaved product. Cleavage can proceed either by aminolysis or by hydrolysis; however, both require cyanylation (G. R. Jacobson et al., 1973; Wu & Watson, 1997). Therefore, we first optimized the conditions for aminolysis, keeping cyanylation conditions constant. In this case, variations in the cleaved/uncleaved ratio indicate variations in cleavage efficiency. Then, we kept highly efficient cleavage conditions constant and varied the conditions of cyanylation. In this case, variations in the cleaved/uncleaved ratio indicate variantions in cyanylation efficiency.

Initial optimization of aminolysis conditions (**Figure 4**) confirmed that the reaction proceeded somewhat better at pH 9, while prolonged incubation had limited effect. The curvature of the bands (“smiling”) near the edge of the gel in Figure 4 is a frequently observed pattern in SDS-PAGE, but it preserves the migration of bands within a lane relative to each other. In all cases, we used the relative intensities of the bands for quantification only for bands within one lane.

**Figure 4.**
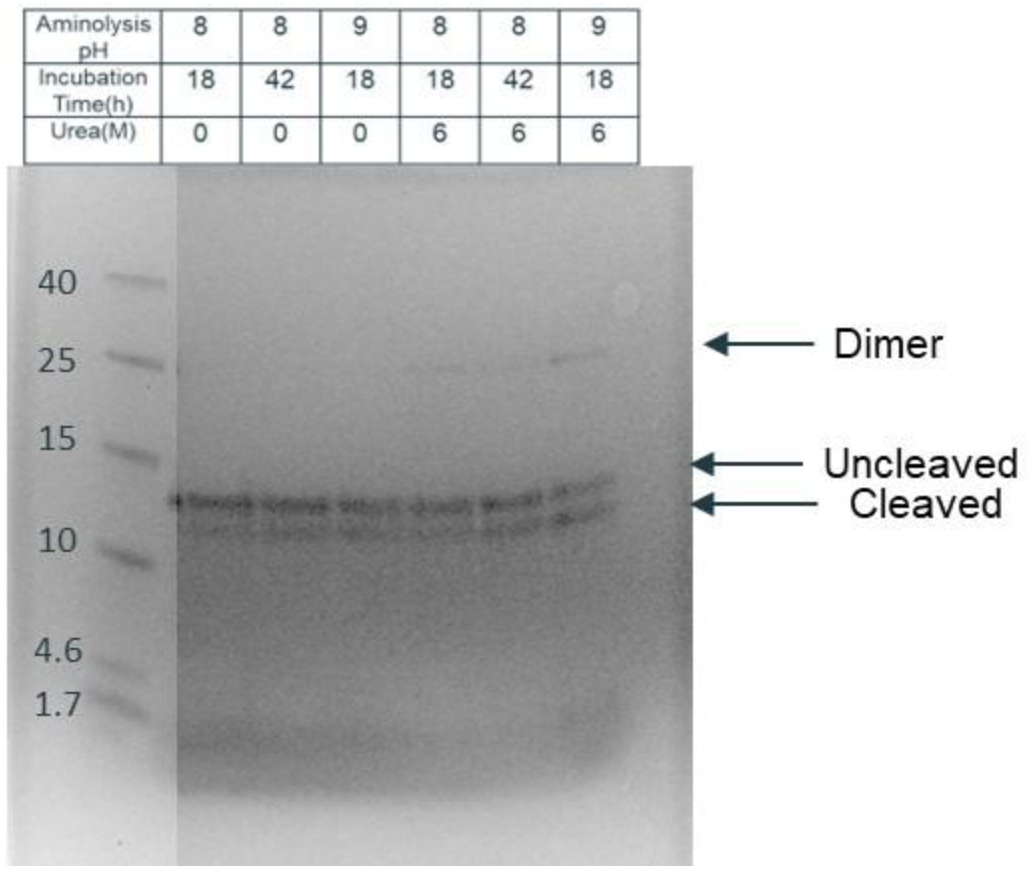
Effect of aminolysis pH and incubation time on cleavage efficiency. Longer aminolysis slightly improved the yield. Aminolysis at pH 9 had higher yield than at pH 8. Higher pH values were not assayed due to competition from hydrolytic cleavage.

We next varied the conditions of the cyanylation step, keeping the pH 9 overnight aminolysis as a constant. We investigated whether urea could improve the reaction rate by increasing structural accessibility of the unique Cys residue (**Figure 5**). Based on cleavage ratios, urea was not necessary for the cyanylation of this recombinant precursor protein, nor did protein concentration have any effect, except that high protein concentration led to aggregation. Next, we optimized the pH, temperature of cyanylation, and concentration of CDAP (**Figure 6**). Incubation at room temperature improved the reaction compared to 4 °C. Increasing CDAP concentration from our initial protocol did not have any effect (all concentrations tested in this figure were much higher than the protein concentration). In fact, very high [CDAP] (50 mM) apparently inhibited the reaction, although we cannot rule out protein aggregation due to exposure to such a high concentration of acetonitrile, the solvent for CDAP. However, we did observe formation of dimers by SDS-PAGE (**Figure 6**). We hypothesized that the free thiol group of reduced Cys could attack the cyanylated Cys residue, leading to formation of a disulfide bond and loss of cyanylation. Reducing these dimers might therefore give the Cys residues another chance to be cyanylated. Indeed, low concentrations of the reducing agent TCEP improved reaction efficiency (**Figure 7**). Conversely, we sought to determine the minimum concentration of CDAP needed for the reaction (**Figure 8**). The minimum concentration without changing the reaction rate was 100µM, or ∼3.5 equivalents.

**Figure 5.**
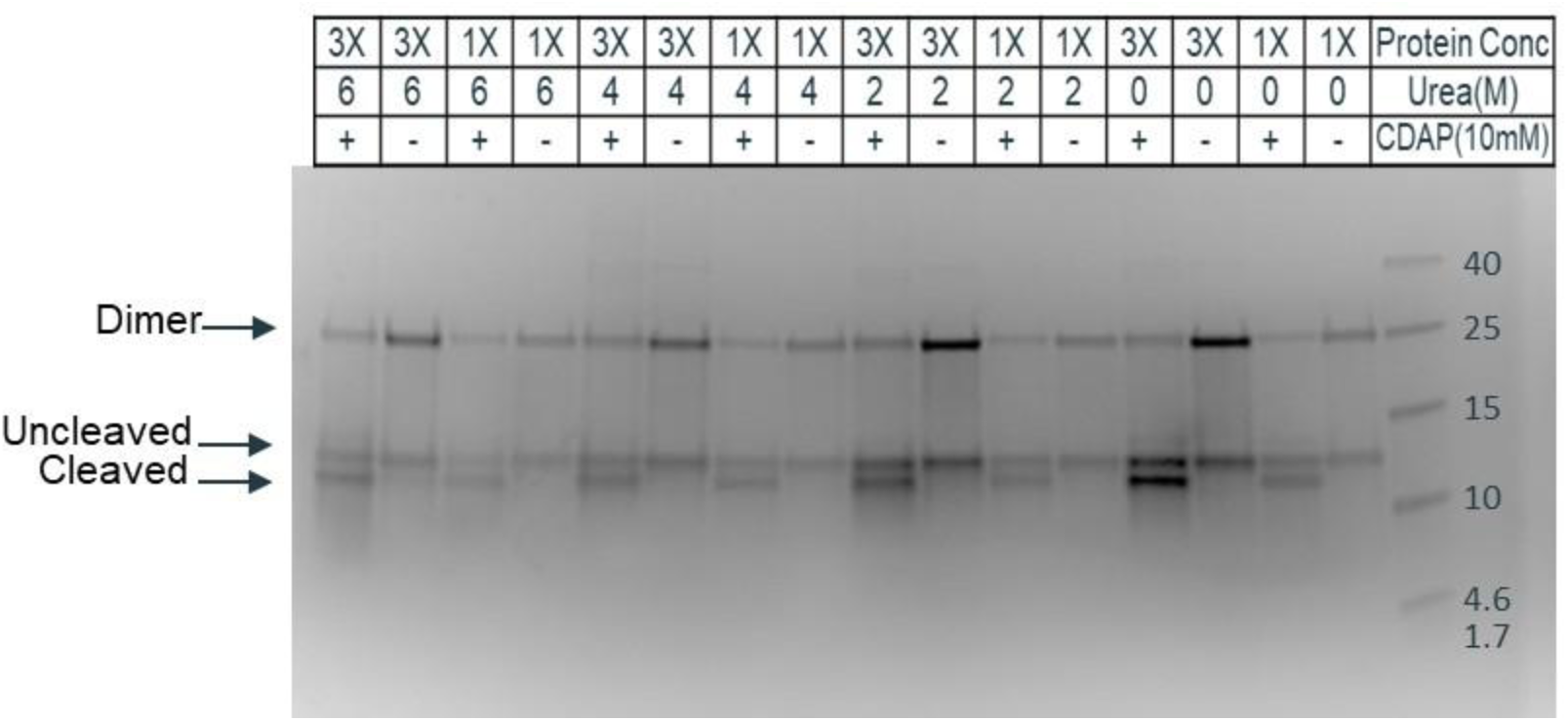
Effect of protein concentration and urea in the reaction. The protein concentration and Urea did not change the reaction significantly, and the rate of all of them remained almost 50%. The 3X corresponds to 27.9 µM and 1X corresponds to 9.3 µM of engineered protein concentration.

**Figure 6.**
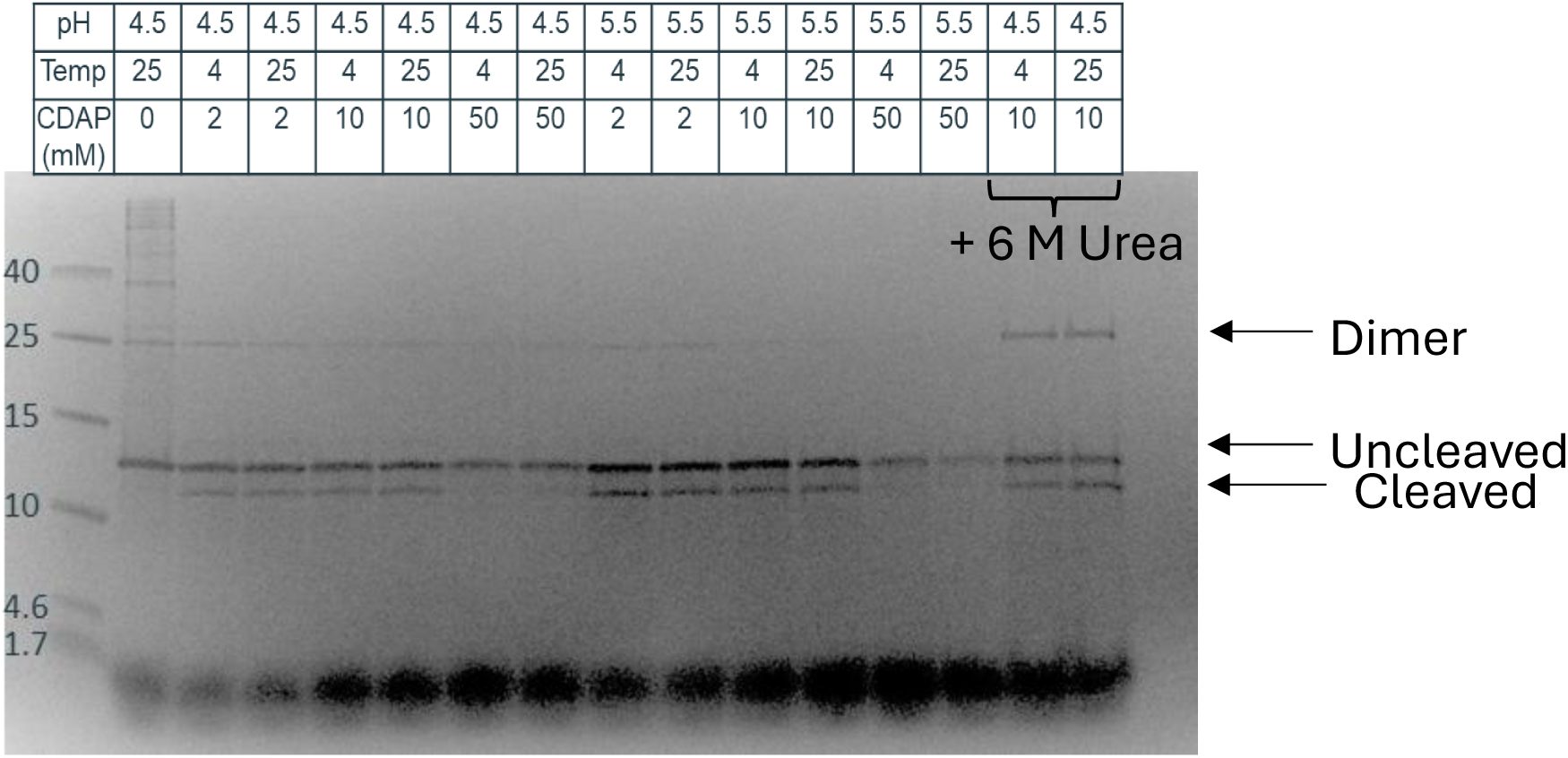
Effects of pH, temperature, and concentration of CDAP on the cyanylation reaction. Cyanylation at room temperature was higher than at 4 °C. Increasing CDAP concentration did not improve the reaction rate. The last two lanes are from reactions containing 6 M urea.

**Figure 7.**
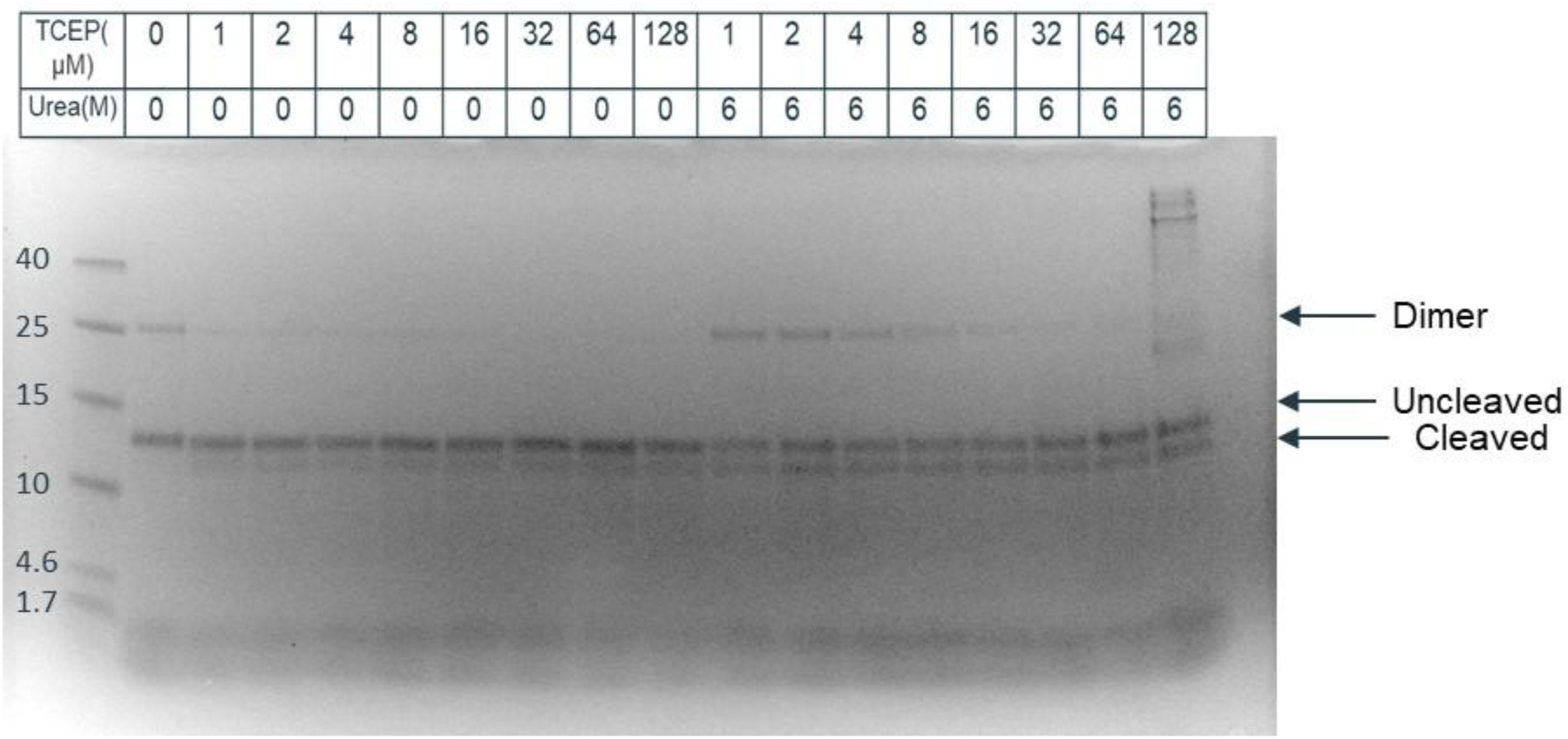
Effect of reducing agent TCEP on cyanylation efficiency in the presence and absence of 6M urea. Higher concentration of TCEP in the presence of urea improved the reaction efficiency by preventing dimer formation. In the absence of urea, equimolar concentration of TCEP improved the reaction efficiency.

**Figure 8.**
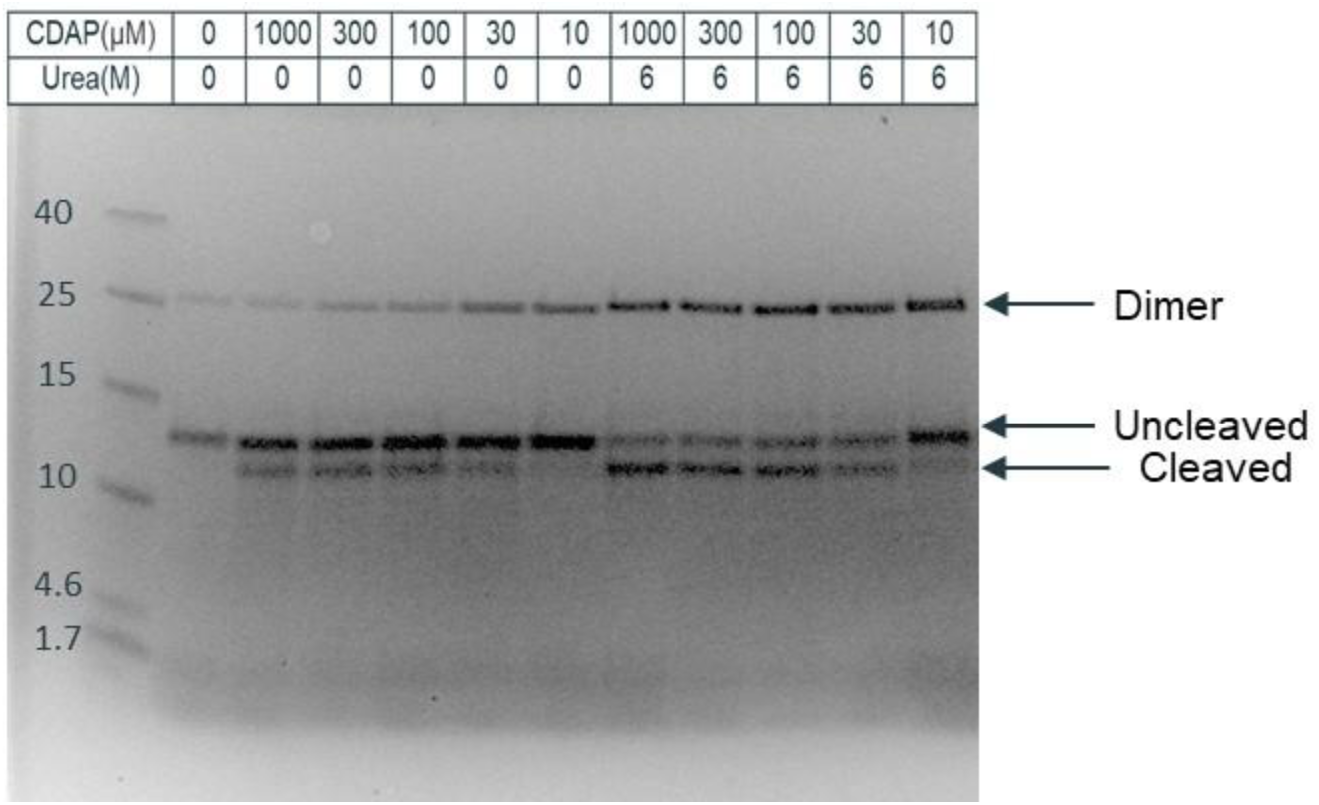
Effect of CDAP concentration in the presence and absence of 6M urea. Decreasing CDAP concentration up to 100 µM did not decrease the reaction efficiency.

### Investigating the effect of peptide composition on cyanylation and cyclization efficiency

To investigate the effect of charge and size of residues in the vicinity of the cysteine on cyanylation and cyclization efficiency, we designed 7 constructs based on the mini-chaperone sequence (**Figure 9**). The charge might induce electrostatic interaction between the first and last amino acids in the peptide, and it may change the reaction rate. The size of amino acids next to cysteine might hinder the attacking N-terminus amino acid. For instance, C1.1 has neutral amino acids, and C1.5 has negatively charged amino acids. These seven constructs were cloned and purified and compared with the original construct. The same protocol was applied to all the eight constructs and cyanylation efficiency was measured by SDS-PAGE. The experiments were done in triplicate and bands were quantified (**Figure 10, Supplementary Figures S1-S4**). Interestingly, C1.5, which has two negatively charged amino acids, had the lowest cyanylation efficiency. This might be due to the efficacy of cyanylation of the cysteine when there are two electron-rich amino acids next to it.

**Figure 9:**
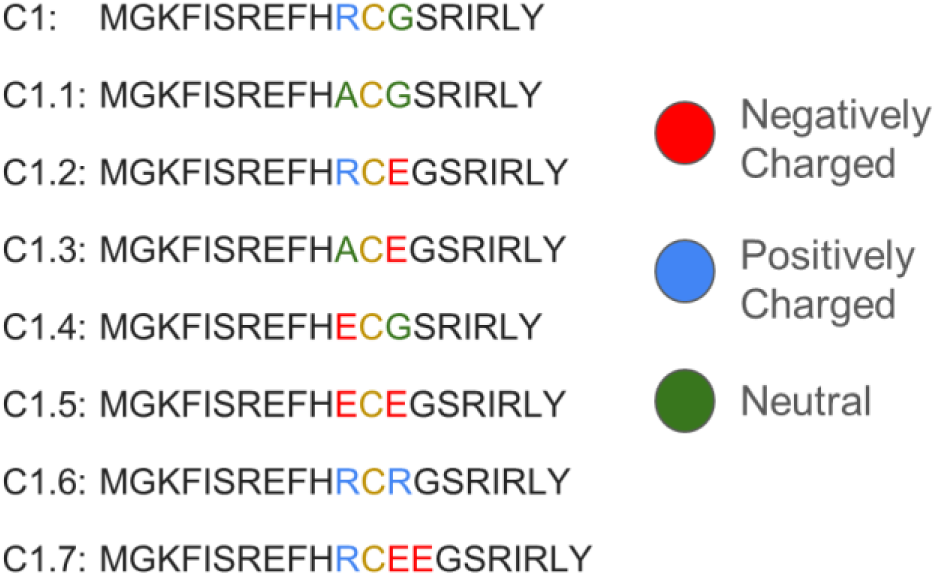
Design of recombinant peptide-protein fusion constructs to investigate the effects of varying the charge of the residues near the unique Cys. The constructs used were as depicted in **Figure 3**. The unique Cys residue that is the site of cyanylation and cleavage is shown in yellow font. The N-terminal Met residue is removed during protein expression, leaving Gly as the N-terminal residue. So, the final sequence of the peptide is “GKFISREFHX,” where X is R, A, or E, as indicated. The residues before and after this Cys were varied as indicated in blue, red, and green font. For simplicity, only the first few residues of the carrier domain (RIRLY) are shown.

**Figure 10.**
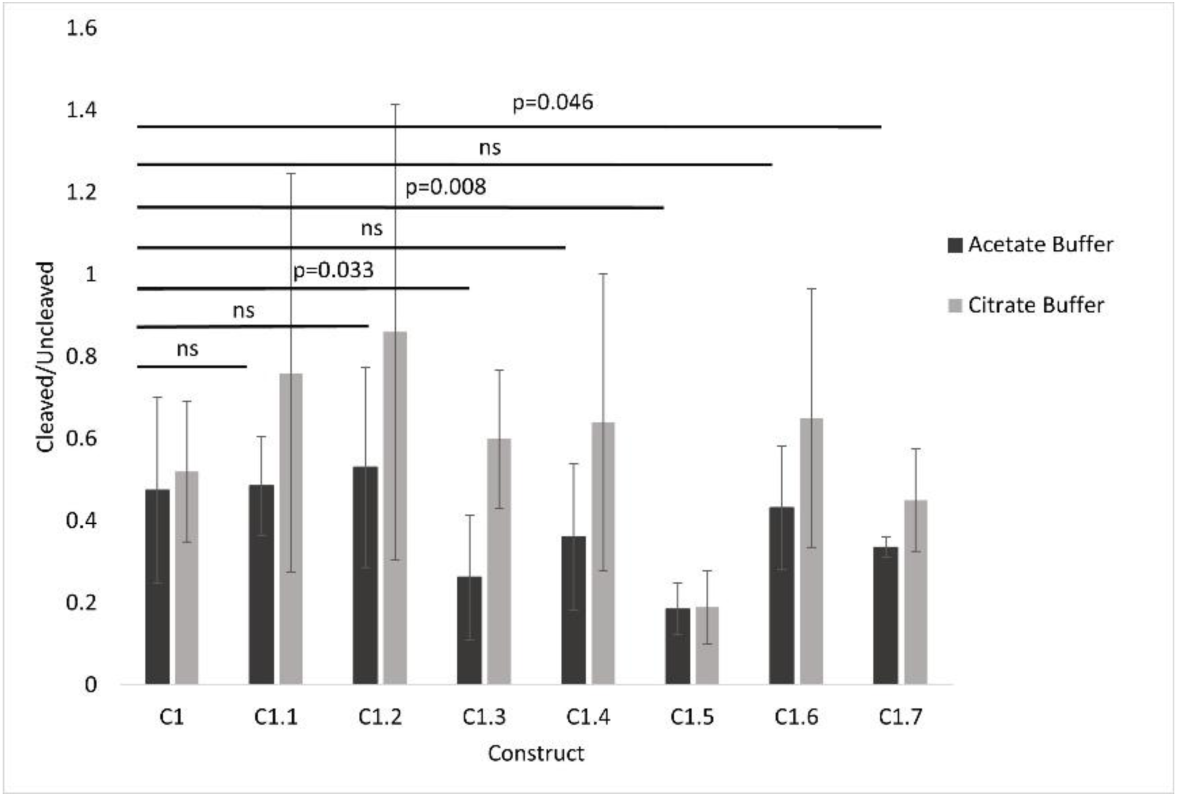
Effect of sequence of peptide on cyanylation efficiency. Rate of cleaved to un-cleaved by using Citrate buffer and Acetate buffer for decreasing pH in the cyanylation step. Data represent mean ± SD from three independent experiments (n = 3). The t-test significance tests between C1 and the other constructs were carried out only for the Acetate approach. Raw data (stained SDS-PAGE gels) are shown in **Figures S1-S4**.

To better understand the effects of the peptide and carrier sequence on cyclization efficiency, we analyzed the reaction of our eight variants by mass spectrometry and assigned ratios based on MS1 ion intensities. The results showed relatively modest effects of the sequence on small differences in terms of produced peptides as the result of the sequence (**Figure 11**). For instance, C1.7 had lower cyclic peptide compared to C1 in the first analysis. Furthermore, by using another analysis approach, we found that C1.3 had lower cyclic peptide products compared to C1.2 and C1.6. Finally, we detected Citrate-conjugated and Acetate-conjugated peptides, respectively, when these buffers were used to change pH during the reaction. However, citrate appeared to interfere with the circularization reaction to a greater extent by blocking the N-terminus. Therefore, we chose Acetate buffer for future scale-up experiments.

**Figure 11.**
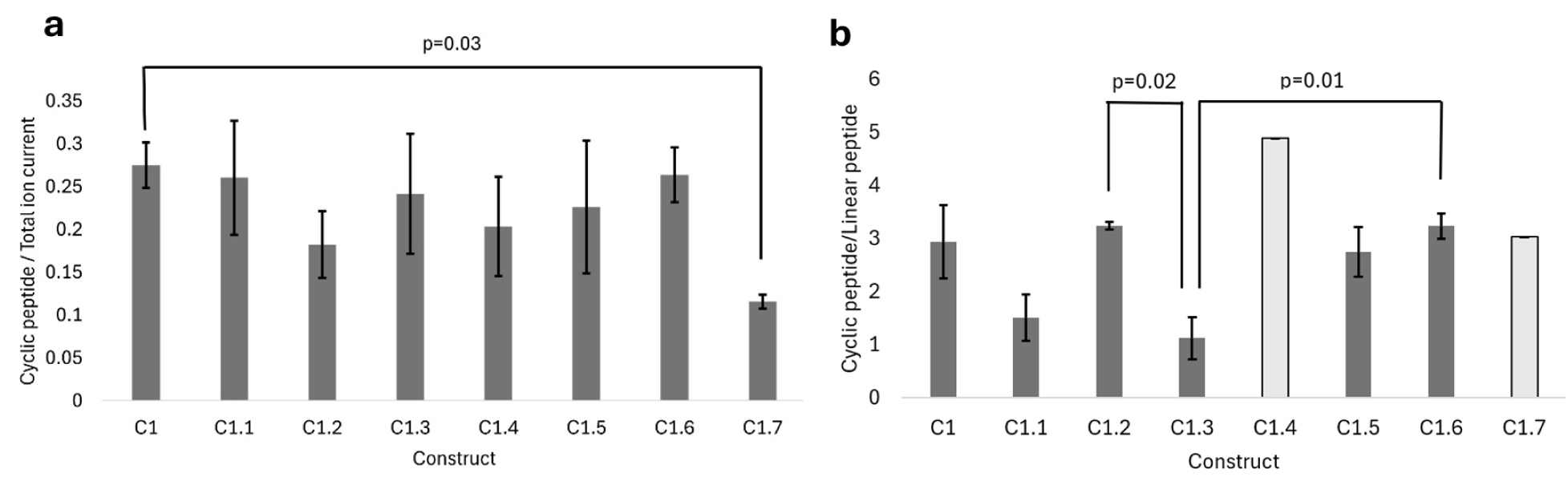
Effect of sequence of peptide on circularization versus linearization. (**a**) Ratio of cyclic peptide ion current to total ion current (sum of linear and cyclic, unmodified and acetylated forms). The only statistically significant difference was observed between C1 and C1.7 (**b**) Ratio of cyclic peptide ion current to linear peptide ion current. For construct C1.4 and C1.7, linear peptide was detected in only one or two replicates, so no statistics could be carried out. The only statistically significant differences were observed between C1.3 versus C1.2 and C1.6. Data represents SD from three independent experiments (n = 3) for the acetate approach.

### Purification of cyclic peptides using HPLC

We next used preparative HPLC to separate cyclic peptides from the mixture. We chose C1.3 for this experiment and prepared 5mL reaction for this purpose. Peptides were purified using RP-HPLC on a C18 column with a linear acetonitrile gradient (**Figure 12** and **Figure S7**) as discussed in the Methods section. The linear and cyclic peptide products eluted at approx. 35% and 40% Acetonitrile concentration, respectively. The cyclic peptide is more hydrophobic than the linear because the linear peptide has hydrophilic amine and carboxyl groups on its termini, while the cyclic peptide lacks termini. In addition, under the acidic HPLC conditions (0.1% formic acid) the linear peptide has an extra positive charge, on the N-terminus, compared to the cyclic. The molecular weights of the linear and cyclic products were confirmed by direct-injection electrospray mass spectrometry on a Bruker Impact II high-resolution instrument.

**Figure 12.**
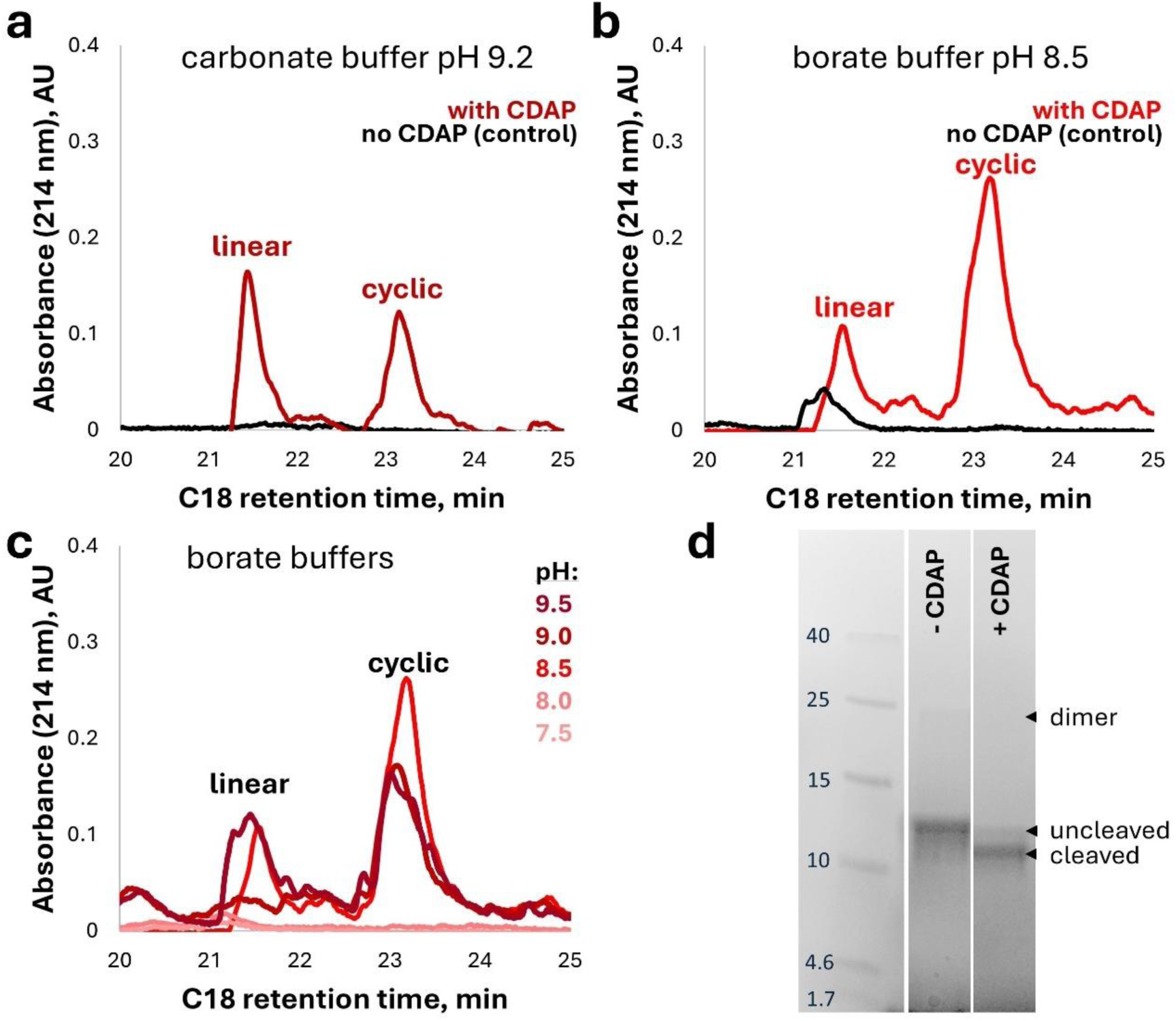
Optimized cyclic peptide production. (a – c) HPLC chromatograms of the linear and cyclic forms of the GKFISREFHA peptide generated from construct 1.3 resolved on a Supelco C18 4.6×15 mm, 5um column with an acetonitrile gradient with 0.1% formic acid. All samples were cyanylated identically in pH 4.5 acetate buffer, except the non-cyanylated control. All samples were desalted after cyanylation into the respective buffers for aminolysis: (a) pH 9.2 carbonate buffer; (b) pH 8.5 borate buffer; (c) borate buffers at pH 7.5 or 8.0 did not facilitate detectable aminolysis, while those at pH 9.0 or 9.5 produced less cyclic and more linear products, as expected, since higher pH favors hydrolysis over aminolysis in aqueous buffers.

**Figure 12** shows linear and cyclic peptide production by the larger-scale method with desalting. Removal of the acetate and DMAP results in cleaner products well resolved by HPLC on a C18 column with an acetonitrile gradient. The buffer salt and the pH are both important at the aminolysis step. Thus, aminolysis in pH 9.2 carbonate buffer (**Figure 12a**) resulted in a higher yield of linear product compared to cyclic, while aminolysis in pH 8.5 borate buffer (**Figure 12b**) greatly favored the cyclic product. In general, higher pH is expected to favor the linear product because it is closer to the pKa of water, thus favoring hydrolysis over aminolysis. Indeed, we observed higher linear/cyclic ratios for borate buffer at pH 9.0 or 9.5 compared to pH 8.5; however, very little of either product was detectable at pH 7.5 or pH 8.0 (**Figure 12c**). Overall yields of aminolysis were much higher at pH 8.5-9.5 than at pH 7.5 or 8.0 (**Figure S6**).

Full HPLC chromatograms, including the peak for the carrier protein, are shown in **Figure S7**. The starting concentration of precursor proteins was identical in all these samples, but some conditions showed a reduction in the intensity of the carrier protein peak, compared to the top-performing condition (borate pH 8.5). We attribute this to aggregation of the carrier protein under the other aminolysis conditions tested here. We therefore estimated peptide yield based on the pH 8.5 condition, where the carrier protein peak is maximally intense. The yield was estimated by determining the absolute peptide/protein concentration in the fractions of the three major HPLC peaks (linear, cyclic, and carrier protein) using the bicinchoninic acid (BCA) test. The entirety of each peak was collected, dried, resuspended in 100 μl of water, and quantified. This test reported 1245.5 μg/ml of the carrier protein, 28.0 μg/ml of the linear peptide, and 108.1 μg/ml of the cyclic peptide, which we corrected down to 108.1 x 9/10 = 97.3 μg/ml because the cyclic 10-mer has 10 peptide bonds instead of the 9 for the linear. The molecular weight of the cleaved carrier protein is 11,517 Da., and the mass of the linear peptide is 1190 Da. We used the same mass for the cyclic peptide, since the BCA assay counts only the number of peptide bonds. In this way, we arrived at the molar percent yield of 11% for the linear and 68% for the cyclic peptide product, for an overall aminolysis yield of ∼78%. Consistent with this estimate, the SDS-PAGE gel in **Figure 12d** shows that aminolytic cleavage of the precursor protein proceeded in very good yield under these optimized conditions. **Figure S8** shows direct-injection electrospray mass spectra of the linear and cyclic peptide peaks, confirming their predominant m/z patterns as expected. Minor unassigned contaminants are present, but their levels relative to the expected products cannot be accurately estimated by mass spectrometry due to likely ionizability differences. The sharp HPLC peaks in **Figure 12a,b** support good purity of both products.

To more fully confirm that the cyclic peptide product was, in fact, cyclic and not a dehydrated peptide, LC/MS/MS was carried out on a ThermoFisher Orbitrap Eclipse instrument in positive mode with HCD fragmentation. As previously observed for the HγD-crystallin derived peptide in **Figure 2**, both the 3+ and 2+ precursor ions corresponding to the expected m/z of the cyclic product yielded highly complex fragmentation patterns. These patterns were consistent with ring-opening at multiple possible sites, followed by fragmentation predominantly from the C-terminus, leading to a preponderance of B-ions. We analyzed the fragmentation patterns manually using Thermo Xcalibur QualBrowser, setting empirical thresholds for MS/MS peak intensities and assigning them by inspection to the set of all possible circular permutants of the “GKFISREFHA” sequence. Our findings are summarized in **Figure 13**. Similarly to **Figure 2**, ring-opening appeared to proceed preferentially within two peptide bonds of the Lys residue. Some of the MS/MS ions assigned in **Figure 13** could have been produced instead as Y-ion fragments of other assigned MS/MS ions; however, we assigned them in these series by parsimony and, to some extent, by similarity of ion current intensities. These choices do not affect the basic conclusion of the assignment: the cyclic peptide was successfully produced, and it opened stochastically at multiple places during MS/MS fragmentation. Details of fragmentation pattern assignment are included in **Supplementary File 1**.

**Figure 13:**
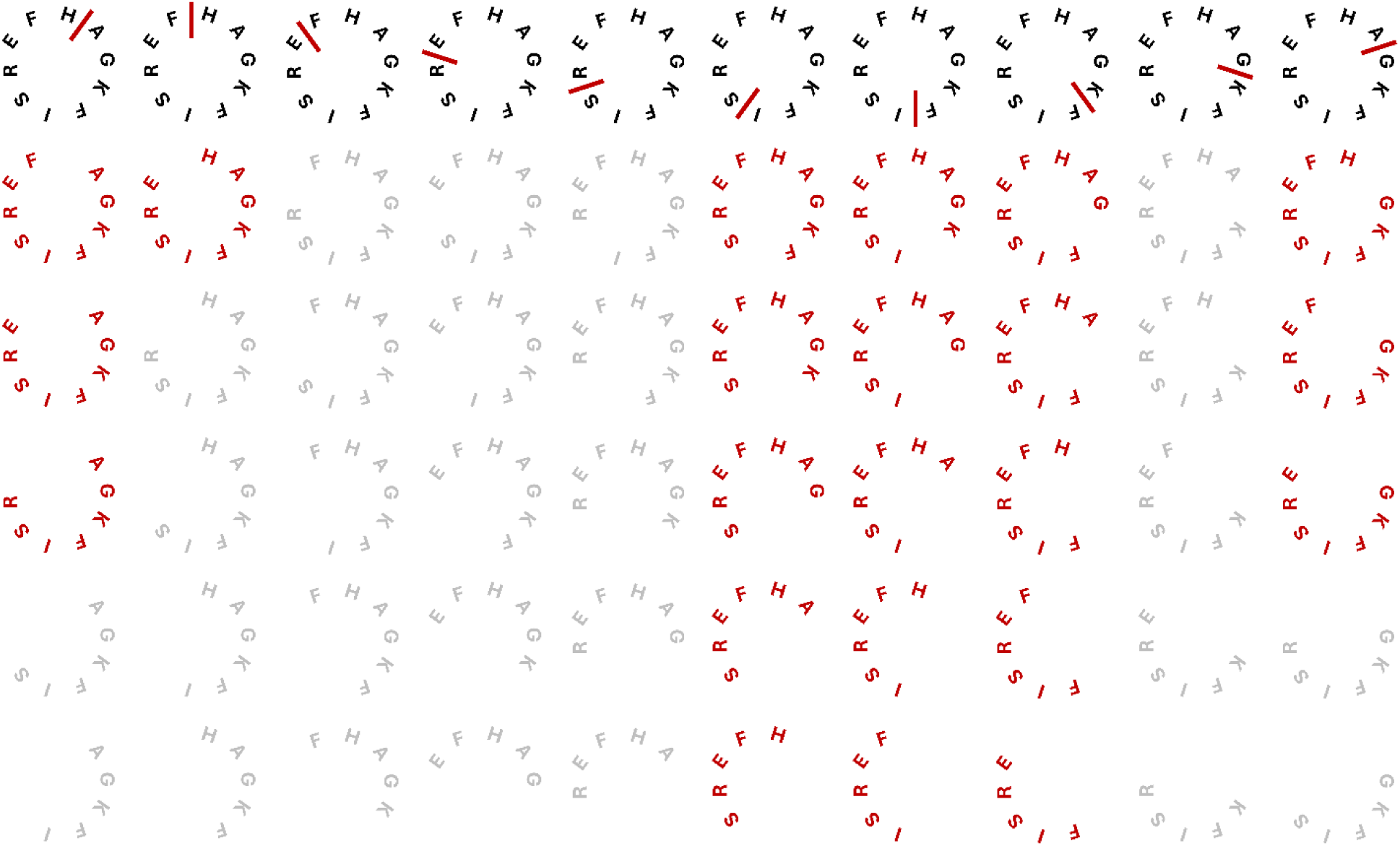
Summary schematic of the most prominent MS/MS fragmentation patterns of the cyclic peptide product. (Chromatogram is shown in **Figure S5**). All possible sites of ring opening are shown as red bars in the top row, and all MS/MS ions with at least five residues and above-threshold ion intensities are shown in red. Fragments shown in gray were theoretically possible but not detected here.

Assignments of the fragmentation patterns in **Figure 13** were made primarily from the 2+ charged precursor at m/z = 587.31, which is an exact match for the precursor expected for the unmodified cyclic peptide. Both fragment ions and neutral loss ions were assigned. Some additional assignments were made from the 3+ precursor at m/z = 391.69, which is close but not exact relative to the expected 391.88 (which was nonetheless observed in this precursor’s MS/MS spectra). We also examined MS/MS spectra for the expected precursor ions for linear, linear-acetylated, and cyclic-acetylated products. All MS/MS m/z tables obtained directly from raw MS/MS data, as well as our assignments, are included in **Supplementary File 1**. Aside from the 2+ charged precursor at m/z = 587.31, no other exact-match precursors were auto-selected for MS/MS. However, we were able to assign traces of the other reaction products by examining the next-closest MS1 precursor m/z’s. Thus, the 3+ charged precursor at m/z = 397.99 was close to the expected m/z of 397.88 at z = 3 for the linear peptide product. MS/MS spectra of this precursor revealed several fragments assignable to fragmentation of the linear product from the N-terminus, in the pattern GK|F|I|S|REFHA, where | indicates observed bond cleavage. Traces of the linear product were expected due to the close spacing of the two HPLC peaks (**Figure 12**) and the high sensitivity of mass spectrometry. Even these MS/MS spectra, however, were dominated by peaks assignable only to the cyclic product, due to multiple circular permutations of the GKFISREFHA sequence. Lys acetylation produces a 42.01 Da. shift in the peptide mass. From the 2+ charged precursor at m/z = 608.13 (vs. 608.32 expected for the acetylated cyclic product), we were able to identify 10 fragment ions consistent with four distinct opening sites of the cyclic peptide acetylated at the Lys residue. The four ring opening sites observed for the acetylated product were A|G, I|S, F|H, and H|A. All four were among the six opening sites observed for the unmodified product in **Figure 13**. It is noteworthy that our cyclic product can only be acetylated at the Lys, since it lacks a free N-terminal amine. Accordingly, we did not observe any B-ions from the K|F ring-opening site (which was detected confidently for the unmodified product), as any C-terminal fragmentation of this ring-opening product would lead to loss of the acetyl-Lys.

## Discussion

The ease and speed of our protocol makes it useful for both small-scale and large-scale production of seamlessly head-to-tail cyclized peptides with only basic equipment and expertise. This method also facilitates quick substitution of amino acids without substantial modification of the protocol; this enables quick in-house production and engineering of cyclic peptides. Furthermore, production of cyclic peptides from recombinant proteins reduces the release of toxic waste compared to solid-phase synthesis. Cyanylation-aminolysis chemistry occurs readily and specifically in water under mild conditions and can therefore be considered an example of “Click” chemistry (Kolb et al., 2001). In contrast, production of peptides by solid-phase synthesis requires using large amounts of environmentally problematic solvents (Martin et al., 2020). Moreover, engineering cyclic peptides from natural amino acids for therapeutic targets is promising since it does not produce toxic byproducts after degradation inside the human body. Additionally, production of peptide from genetically-encoded material will accelerate sharing these peptides between laboratories since the DNA sequences and plasmids can be easily shared.

Production of cyclic peptides recombinantly limits incorporation of non-natural amino acids into cyclic peptides since non-natural amino acids require their own tRNA (Jin et al., 2019). Therefore, it is best suited for production of compounds that mimic naturally occurring ribosomally synthesized cyclic peptides, known as orbitides, bacteriocins, etc. However, recombinant precursor proteins could also be produced via cell-free expression, which allows for incorporation of a wider range of non-natural amino acids (Gao et al., 2019), though it also increases the cost.

We have demonstrated here that seamlessly head-to-tail cyclized peptides can be excised from proteins via cyanylation-aminolysis. These peptides resemble naturally-occurring orbitides in being cyclized only with peptide bonds, being composed of only L-amino acids, and lacking disulfides. Our objective was only to prove feasibility, so we did not rigorously calculate the final purity and yield of the products. However, we can estimate the overall yield from the apparent yields of the two steps of the reaction. Efficiency of the small scale cyanylation method ranged from ∼20-50% (**Figure 10**) in acetate buffer, with potentially higher efficiencies in citrate buffer but at the cost of high variability and adduct formation. We assigned four reaction products based on MS1 precursor ions: linear, cyclic, and acetylated versions of either. Yields of unmodified cyclic products relative to all assignable peaks averaged ∼20% (**Figure 11a**), for an overall estimated yield of 4-10% for the small-scale method. This would make the yields from our approach comparable to those of traditional cyclization reactions used in solid-phase synthesis but likely underestimates the true yield because cyclic peptides lack the N-terminal free amine group and are therefore less ionizable than linear products of the same sequence.

The optimized large-scale method was more efficient, likely due to the benefit of desalting after the cyanylation step. Without desalting, a significant proportion of the product was linear and acetylated. We hypothesize that N-terminal acetylation competes with intramolecular aminolysis during the high-pH incubation, and that removal of the acetate buffer by desalting prevents this. Combined with the optimized borate-buffered aminolysis condition, we achieved 68% yield of c(GKFISREHA) cyclic peptide product. This is comparable to enzymatic cyclization methods.

Moreover, the observed ratios of cyclic to linear product ions varied based on the residues in the immediate vicinity of the Cys residue (**Figure 11b**). The observed variation suggests that small residues near the Cys promote cyclization, perhaps by ensuring good accessibility of the peptide bond to be transferred. Conversely, negatively charged (Glu) residues, particularly after the Cys, may inhibit cyclization and favor the linear hydrolysis product. Future experiments should test our method on a wider variety of peptide lengths and sequence compositions to determine how those parameters affect the overall yield and the linear/cyclic ratio. In general, we predict that longer peptides will have higher linear/cyclic ratios, since the termini are further apart; however, if the peptide folds into a structure that brings the N-terminus closer to the cyanylated Cys residue, then the length of the peptide may not matter.

Producing comparable quantities of linear and cyclic peptides is a useful feature at the research stage, since in many applications it is of interest to compare the performance of the linear and cyclic versions of the same peptide sequence. In this case, both are produced in one pot. For optimal scale-up, however, it will be important to further improve cyanylation chemistry, rigorously quantify the overall reaction yields, further optimize the aminolysis conditions, and engineer improved carrier proteins for maximum expression, solubility, and suitability for the reaction. It will also be important to investigate how a peptide’s length and structure affect cyclization yields. Our cyclic 17-mer (**Figure 2b**) and 10-mers (**Figure 11**) were produced with comparable cyclic:linear ratios; however, the 17-mer’s β-hairpin propensity may have facilitated this. Cyclization methods in general may favor hairpin-forming sequences due to greater average proximity of their termini.

Generation of acetylated side products was a surprise given our reaction composition. Acetylation of the N-terminus occurs, but MS/MS suggests some acetylation of Lys, as well. Non-enzymatic pathways of Lys acetylation are known (Baldensperger & Glomb, 2021), but they involve compounds such as acetyl phosphate. This was unlikely to form in our reaction, since all proteins were desalted out of phosphate buffer by size exclusion. It remains to be explored whether reaction conditions that promote aminolysis also promote Lys acetylation. However, removal of acetate by desalting mitigated the acetylation issue.

An intriguing future possibility opened by our approach is to generate large libraries of seamlessly cyclic peptides at very low cost by excising them directly from natural proteomes. Natural proteome-derived linear peptide libraries have already been widely used, primarily for characterizing specificities of novel proteases (Biniossek et al., 2016; Schilling & Overall, 2008; Tanco et al., 2013; Tucher et al., 2014) or other peptide modification enzymes and reagents (Bridge & Weeks, 2023; Van Damme et al., 2011; Weeks & Wells, 2018). In at least one case, a proteome-derived linear peptide library was directly used to screen for binders of protein targets (Maaty & Weis, 2016). Applying our present strategy to this problem promises highly scalable, inexpensive generation of peptide macrocycles of a wide variety of sizes and structures, in libraries whose sequence composition is derived from natural proteomes and may therefore be more likely to yield biologically available and active initial hits. (The hits may subsequently be optimized using focused, designed libraries.) By choosing which protein or proteomic samples to use as starting material and whether and how to digest them beforehand, researchers have a meaningful degree of control over the final composition of the cyclic peptide library thus produced. Initial screening for biological activity may then be analogous to the long-established drug discovery approach of screening plant extracts for desired bioactivity, then fractionating the extracts to isolate the active ingredient. In essence, we propose a way to “extract” cyclic peptides not only from the sap of trees but from arbitrarily chosen mixtures of natural protein feedstocks.

## Methods

### Protein expression and purification

WT HγD Protein expression and purification were performed as previously described by Serebryany et al. (2015).

The protein-fusion constructs were cloned, expressed and purified as follows. The backbone of pET16b plasmid containing sGFP was linearized by PCR. The gene of sGFP was omitted and synthesized gene containing homology region to the backbone was used for in-vivo assembly. The bacterial colonies were plated on Ampicillin plates, and their plasmid were sequenced by GENEWIZ. pET16b plasmids containing the genes of variants were transformed into BL21(DE3) and selected with 100µg/mL Ampicillin. Single colonies were used to inoculate 10mL TB (Casein digest peptone 12g/L, Yeast extract 24g/L, Dipotassium Phosphate 9.5g/L, Monopotassium Phosphate 2.2g/L) in 50 mL conical flask which were cultured overnight at 37 C, 220 rpm overnight. Overnight cultures were used to inoculate main cultures (1 to 100). Cultures were grown under 37 °C and 220 rpm until reaching OD ≈ 0.6. Then the expression was induced by adding IPTG to the final concentration of 1mM. The cultures were induced overnight at 16 °C, and harvested the following by centrifugation at 4000 g for 40 minutes at 4 °C. Bacterial pellets were resuspended in Binding Buffer (50mM sodium phosphate, 300mM sodium chloride, 10mM imidazole; pH 7.4) and protease inhibitor (Thermo Scientific Pierce mini tablets, one tablet per 600mL culture) was added. The bacterial cultures were lysed by 30 minutes incubation with lysozyme (0.1mg per 600mL of culture) and sonication (Fisherbrand Model 120 Sonic Dismembrator with 1/8 inch probe, 70% Amplitude, 2s pulse, 2s rest for 5 minutes total sonication time) on ice. Next, the suspension was centrifugated (40000 g, 30 minutes, 4 °C) to separate bacterial debris from soluble proteins. The supernatant was injected into HisPur Cobalt chromatography (Thermo Fisher) column and was purified by Elution Buffer (50mM sodium phosphate, 300mM sodium chloride, 150mM imidazole; pH 7.4). The eluted proteins after this step were incubated with 5mM TCEP for 1h to reduce any possible disulfide bonds. The purified proteins from affinity chromatography were applied to size exclusion chromatography (Superdex75). The proteins then were purified in size-exclusion buffer (10mM PIPES, 50mM NaCl, pH 6.7). Purified proteins were then aliquoted and stored at −70.

### Small-scale cyanylation and aminolysis

For the cyanylation step, the pH was decreased to 4.5 either by adding Citrate buffer (1M, pH 4) or Acetate buffer (1M, pH 4). Protein concentration was 27 μM. Then CDAP from the stock (400mM, dissolved in acetonitrile; Thermo Fisher Scientific) was added to the reaction (final concentration of 10mM) and incubated at room temperature for two hours. The pH then was increased to 9 by adding Carbonate buffer (1M, pH 9.2) to the reaction for the aminolysis step. The reaction then was incubated at 4 °C for 18 hours. The reaction then was quenched by adding 0.1% Formic Acid.

### Large-scale optimized cyanylation and aminolysis

The pH of 5 mL of protein construct (40 μM) was decreased to 4.5 by adding Acetate buffer (from 1M stock at pH 4). Then CDAP from the stock (400mM, dissolved in acetonitrile; Thermo Fisher Scientific) was added to the reaction (final concentration of 10mM) and incubated at room temperature for two hours. 32 μM TCEP was included in the reaction. Then the reaction was desalted using HiPrep desalting column (Sephadex G-25; Cytiva) into the aminolysis buffer (Borate buffer, 20 mM, pH 8.5, or others as indicated). The reaction then was incubated at 37 °C overnight. The reaction then was quenched by adding formic acid to 0.1% v/v final.

### SDS-PAGE and quantification

The purity of protein purification was assessed by 12% SDS-PAGE using precast BioRad Criterion gels and GelCode Coomassie stain (ThermoFisher). The rate of cleavage of the constructs was assessed by running 16% SDS-PAGE and fixing the gels before staining according to the manufacturer’s protocol. Quantification of bands was done by GelAnalyzer software.

### 1D mass spectrometry for measuring cyclization

The samples after completion of the reaction were quenched by 0.1% formic acid. About 40µL of each sample at 20µM concentration was applied to column (Acquity UPLC protein BEH C4 column 2.1 x 50 mm). peptides were separated using a 11 min gradient at 300µL/min by 5%, 6.5 min at 40%, 7 min at 98%, hold for 8.75 min, 9 min at 5%, and hold for 11 min. Sample were injected via an autosampler that was kept at 4°C, and the column was kept at 40°C. Eluting peptides were analyzed by mass spectrometry (Waters Xevo-GS, positive polarity, and acquired over 110-2000 m/z at 0.2 scans/sec). Lockspray data acquired on LeuEnk peptide every 10 sec and 3 scans averaged to apply real-time correction.

### Preparative HPLC

The mixture was filtered by 0.2 µm membrane and purified using reverse-phase high-performance liquid chromatography (RP-HPLC) on a C18 column (150 mm × 4.6 mm, 5 μm particle size, 100 Å pore size; Sigma-Aldrich). The separation was performed on a Teledyne ISCO ACCQPrep HP150 chromatograph, with UV detector set to 214 nm and 254 nm. Mobile phases were: water with 0.1% formic acid (solvent A) and acetonitrile with 0.1% formic acid (solvent B). Cyclic peptides were purified using a linear gradient from 0% to 100% solvent B for 40 minutes at a flow rate of 1.0 mL/min. The column temperature was maintained at 25 °C during the run. Fractions were collected, concentrated and stored at −20°C until further analysis.

### Measurement of peptide and protein amounts

Purified peaks from HPLC were dried and then resuspended in 100 μL water. An aliquot of 25 μL was taken from each peak and quantified using Bicinchoninic Acid (BCA) protein assay kit (G-Biosciences, MO, USA) by adding to 200 μL of working solution and reading absorbance at 562 nm in a 96-well plate. Standard curve was constructed by serial dilution of bovine serum albumin (G-Biosciences, MO, USA). The concentration of protein was calculated from the slope of standard calibration curve.

### 2D mass spectrometry for detecting cyclic peptide fragmentation patterns

Data in **Figure 2d** were collected on a ThermoFisher qExactive HF+ mass spectrometer in positive ion mode with a standard 1h acetonitrile gradient on a NanoFlow C18 column. The sample was extensively desalted using a C18 desalting cartridge prior to loading to remove GdnHCl. MS/MS data summarized in **Figure 13** were collected on a ThermoFisher Orbitrap Eclipse mass spectrometer, with a Waters BEH C18 column and a 1h acetonitrile gradient, in positive ion mode, HCD fragmentation mode, and 15-second dynamic exclusion window to maximize MS/MS spectral quality for highly abundant precursors. All data were analyzed in Thermo XCalibur QualBrowser by inspection against a manually constructed database of circularly permuted peptide variants and the masses of their B-ion fragments (included in **Supplementary File 1**).

## Acknowledgements

The authors are grateful to Drs. Jennifer Liddle and Jeremy Balsbaugh and the CORE^2 Proteomics Facility at the University of Connecticut; Prof. John Haley and the Biological Mass Spectrometry Core Facility at Stony Brook University School of Medicine; and to Dr. Beniam Berhane and the CASDA Mass Spectrometry Core Facility at Stony Brook University for mass spectrometry experiments and advice. This research was funded by the National Institutes of Health, award R00GM141459.

## Notes

### Competing Interest Statement

The authors have declared no competing interest.

