## Supplementary Figures for "Enzyme-free biochemical production of seamlessly N-to-C cyclized peptides from natural or recombinant proteins"

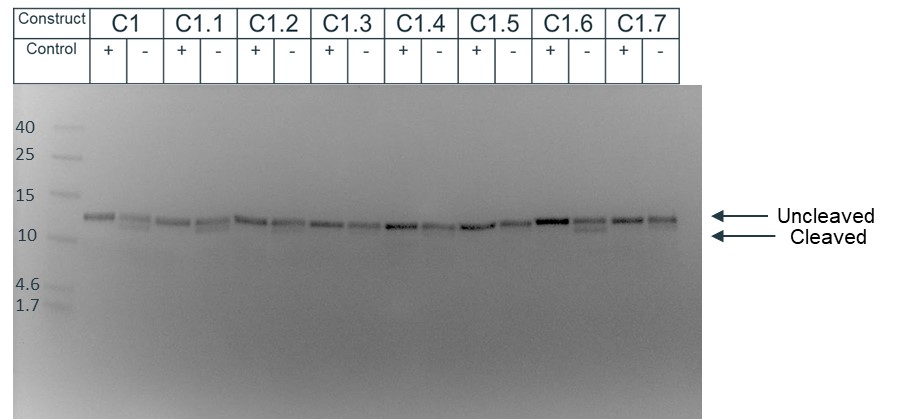

**Figure S1.** **Investigating the effect of peptide composition on cyanylation: First replicate of the citrate buffer cyanylation approach**. Coomassie-stained 16.5% SDS-PAGE gel (Bio-Rad Criterion Tris-Tricine) showing results of backbone cleavage of the eight constructs, as well as the respective control samples. No CDAP was included in the control samples, so they were not cyanylated and consequently not cleaved at the aminolysis step.

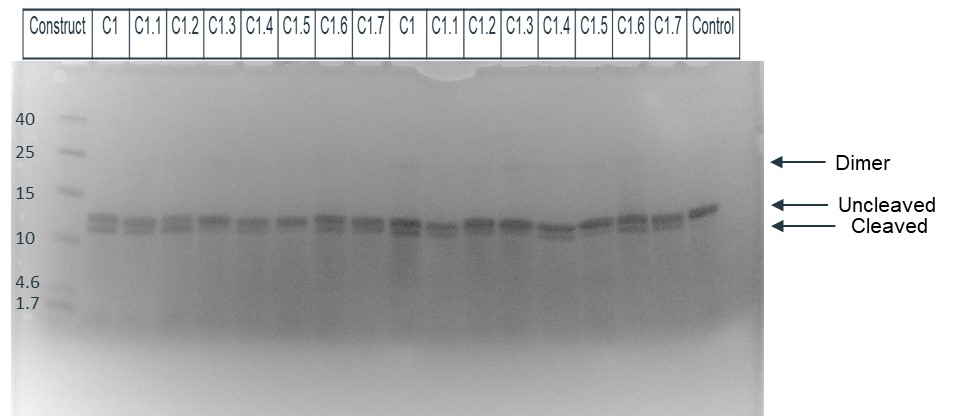

**Figure S2. Investigating the effect of peptide composition on cyanylation: Second and third replicates of the citrate buffer cyanylation approach**. Coomassie-stained 16.5% SDS-PAGE gel (Bio-Rad Criterion Tris-Tricine) showing results of backbone cleavage of the eight constructs, as well as the respective control samples. No CDAP was included in the control sample, so it was not cyanylated and consequently not cleaved at the aminolysis step.

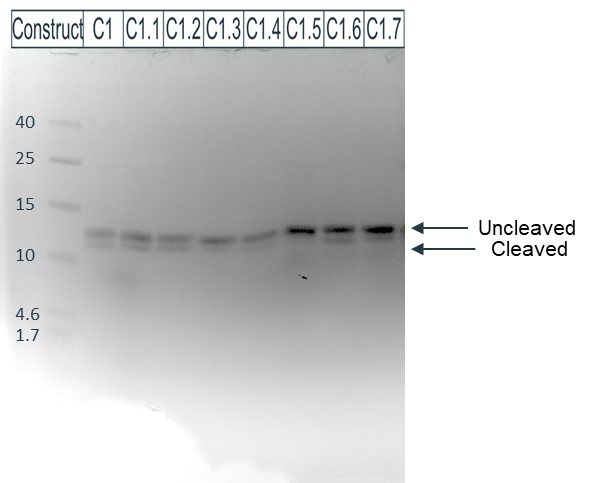

**Figure S3*.* Investigating the effect of peptide composition on cyanylation: First replicate of the acetate buffer cyanylation approach**. Coomassie-stained 16.5% SDS-PAGE gel (Bio-Rad Criterion Tris-Tricine) showing results of backbone cleavage of the eight constructs, as well as the respective control samples.

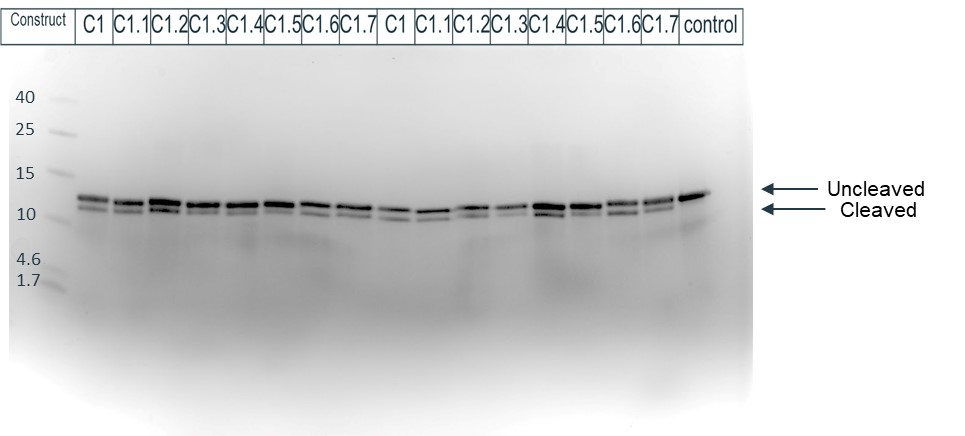

**Figure S4*.* Investigating the effect of peptide composition on cyanylation: Second and third replicates of the acetate buffer cyanylation approach**. Coomassie-stained 16.5% SDS-PAGE gel (Bio-Rad Criterion Tris-Tricine) showing results of backbone cleavage of the eight constructs, as well as the respective control samples. No CDAP was included in the control sample, so it was not cyanylated and consequently not cleaved at the aminolysis step.

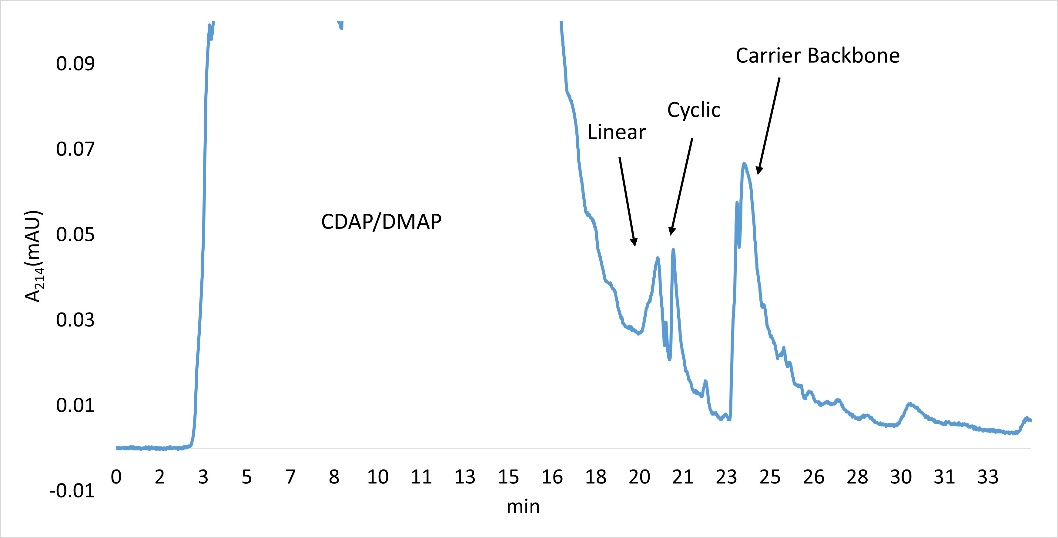

**Figure S5. HPLC chromatogram for the cyclic peptide production reaction without desalting after the cyanylation step.** Cyanylation was carried out in acetate buffer pH 5 for 2 h with 3 molar equivalents of TCEP. Aminolysis was carried out in carbonate buffer pH 9 at 4 °C for 18 h.

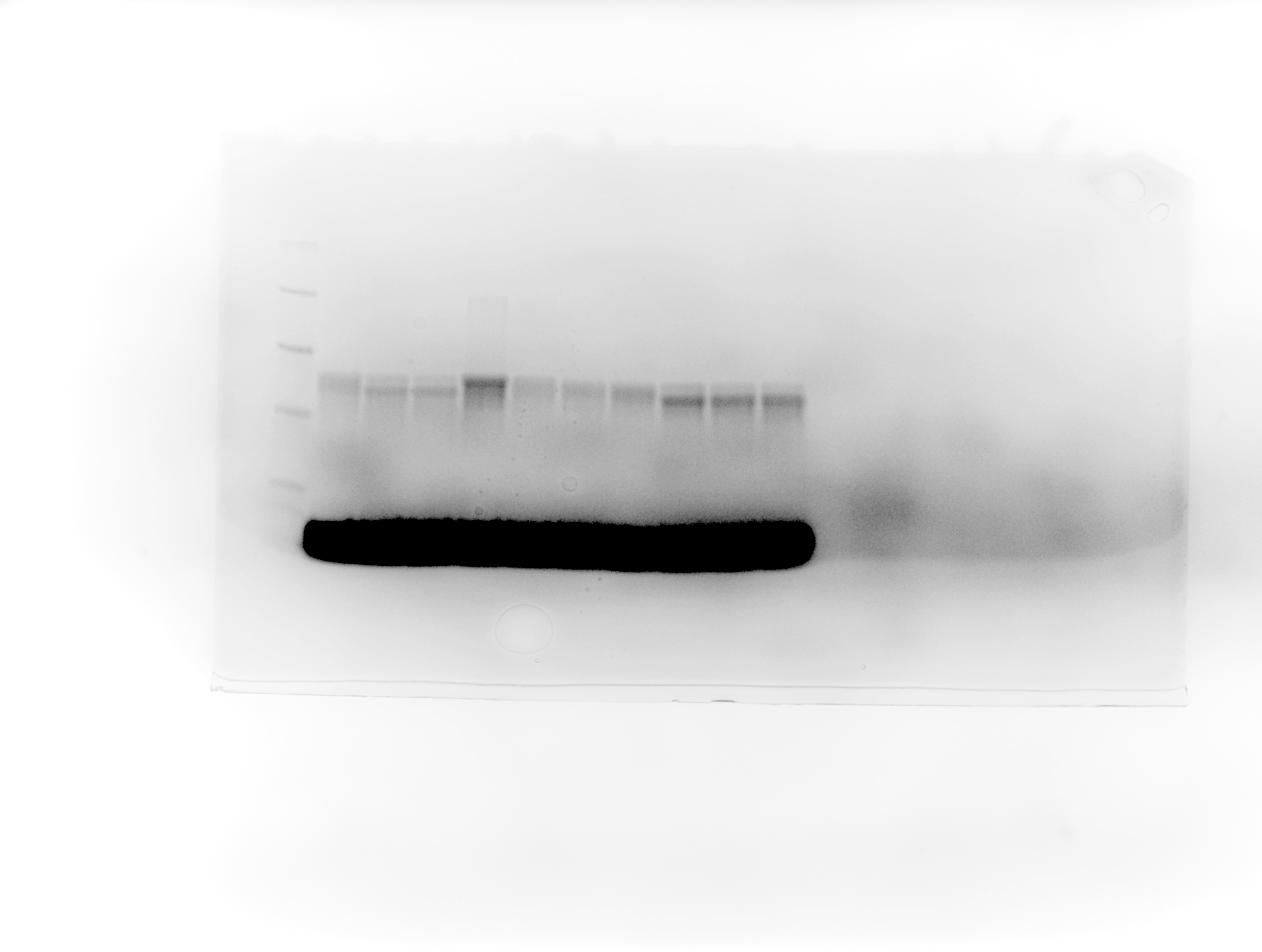

4.6

10

15

25

40

1.7

1

2

3

4

5

6

7

8

9

10

**Figure S6. Optimization of aminolysis in carbonate and borate buffers.** Coomassie-stained 16.5% SDS-PAGE gel (Bio-Rad Criterion Tris-Tricine) showing results of backbone cleavage of construct 1.3, from samples cyanylated identically in pH 4.5 acetate buffer. Buffers and temperatures were varied for the aminolysis step. *Lane 1*: Carbonate buffer (20 mM, pH 9.2), control. *Lane 1*: Carbonate buffer (20 mM, pH 9.2) incubated at 4 °C for 42 hours. *Lane 2*: Carbonate buffer (20 mM, pH 9.2) incubated at 37 °C overnight. *Lane 3*: Borate buffer (20 mM, pH 8.5), control. *Lane 4*: Borate buffer (20 mM, pH 8.5) incubated at 4 °C for 42 hours. *Lane 5*: Borate buffer (20 mM, pH 7.5) incubated at 37 °C overnight. *Lanes 6 and 7*: Borate buffer (20 mM, pH 8) incubated at 37 °C overnight. *Lane 8*: Borate buffer (20 mM, pH 8.5) incubated at 37 °C overnight. *Lane 9*: Borate buffer (20 mM, pH 9) incubated at 37 °C overnight. *Lane 10*: Borate buffer (20 mM, pH 9.5) incubated at 37 °C overnight.

Absorbance (214 nm), AU

| C18 retention time, min | Borate, pH 7.5, 37 °C , Overnight |
| --- | --- |
| 0.050231 | -0.002 |
| 0.0518514 | -0.0021 |
| 0.0680549 | -0.0016 |
| 0.0858788 | -0.002 |
| 0.102082 | -0.0005 |
| 0.118286 | -0.0018 |
| 0.13449 | -0.0023 |
| 0.152313 | -0.0014 |
| 0.168517 | -0.0018 |
| 0.184721 | -0.0031 |
| 0.202544 | -0.0012 |
| 0.220368 | -0.0023 |
| 0.236572 | -0.0015 |
| 0.252775 | -0.0004 |
| 0.270599 | -0.0019 |
| 0.286803 | -0.0002 |
| 0.303006 | -0.0001 |
| 0.31921 | -0.0016 |
| 0.337034 | -0.0016 |
| 0.353237 | -0.0012 |
| 0.369441 | -0.0012 |
| 0.387265 | -0.0013 |
| 0.403468 | -0.0012 |
| 0.419672 | -0.0017 |
| 0.437496 | -0.0008 |
| 0.453699 | -0.0004 |
| 0.469903 | -0.0008 |
| 0.486106 | -0.0012 |
| 0.50393 | -0.0017 |
| 0.520134 | -0.0005 |
| 0.536338 | -0.002 |
| 0.554161 | -0.0023 |
| 0.570365 | -0.0018 |
| 0.586569 | -0.0015 |
| 0.602772 | -0.0017 |
| 0.620596 | -0.0021 |
| 0.636799 | -0.0014 |
| 0.653003 | -0.003 |
| 0.670827 | -0.0029 |
| 0.68703 | -0.0014 |
| 0.703234 | -0.0025 |
| 0.719438 | -0.0026 |
| 0.737262 | -0.0021 |
| 0.753465 | -0.0016 |
| 0.769669 | -0.001 |
| 0.787493 | -0.0027 |
| 0.802076 | -0.0012 |
| 0.818279 | -0.0019 |
| 0.834483 | -0.0019 |
| 0.852307 | -0.0015 |
| 0.86851 | -0.0007 |
| 0.884714 | -0.0025 |
| 0.900917 | -0.0009 |
| 0.918741 | -0.0016 |
| 0.934945 | -0.002 |
| 0.951148 | -0.0012 |
| 0.968972 | -0.0011 |
| 0.985176 | -0.0014 |
| 1.00138 | -0.001 |
| 1.01758 | -0.0007 |
| 1.03541 | -0.0007 |
| 1.05161 | -0.0011 |
| 1.06781 | -0.0022 |
| 1.08564 | -0.0027 |
| 1.10184 | -0.002 |
| 1.11804 | -0.0019 |
| 1.13425 | -0.0025 |
| 1.15207 | -0.0024 |
| 1.16828 | -0.0036 |
| 1.18448 | -0.0024 |
| 1.2023 | -0.0016 |
| 1.21851 | -0.003 |
| 1.23471 | -0.0025 |
| 1.25253 | -0.0033 |
| 1.26874 | -0.003 |
| 1.28494 | -0.0034 |
| 1.30114 | -0.0039 |
| 1.31897 | -0.0048 |
| 1.33517 | -0.0048 |
| 1.35138 | -0.003 |
| 1.3692 | -0.0046 |
| 1.3854 | -0.0037 |
| 1.40161 | -0.0038 |
| 1.41781 | -0.0037 |
| 1.43563 | -0.0048 |
| 1.45184 | -0.0047 |
| 1.46804 | -0.0039 |
| 1.48587 | -0.0042 |
| 1.50207 | -0.0042 |
| 1.51827 | -0.0045 |
| 1.53448 | -0.0041 |
| 1.5523 | -0.0036 |
| 1.5685 | -0.0026 |
| 1.58471 | -0.0015 |
| 1.60253 | -0.0022 |
| 1.61874 | -0.003 |
| 1.63494 | -0.0023 |
| 1.65114 | -0.0027 |
| 1.66897 | -0.0025 |
| 1.68517 | -0.0017 |
| 1.70137 | -0.0015 |
| 1.71758 | -0.002 |
| 1.7354 | -0.0006 |
| 1.7516 | -0.0017 |
| 1.76781 | -0.0008 |
| 1.78563 | -0.0015 |
| 1.80184 | -0.0009 |
| 1.81804 | -0.0011 |
| 1.83424 | -0.0012 |
| 1.85207 | -0.0016 |
| 1.86827 | -0.0011 |
| 1.88447 | 0.0003 |
| 1.9023 | -0.0007 |
| 1.9185 | 0.0011 |
| 1.9347 | 0.0003 |
| 1.95253 | 0.0001 |
| 1.96873 | -0.0005 |
| 1.98493 | 0.0006 |
| 2.00114 | 0.0009 |
| 2.01896 | 0.0006 |
| 2.03517 | 0.0009 |
| 2.05137 | 0.0009 |
| 2.06919 | 0.003 |
| 2.0854 | 0.0022 |
| 2.1016 | 0.0026 |
| 2.1178 | 0.0026 |
| 2.13563 | 0.0026 |
| 2.15183 | 0.004 |
| 2.16804 | 0.005 |
| 2.18586 | 0.0055 |
| 2.20206 | 0.0058 |
| 2.21827 | 0.006 |
| 2.23447 | 0.0078 |
| 2.25229 | 0.0078 |
| 2.2685 | 0.009 |
| 2.2847 | 0.0106 |
| 2.3009 | 0.0101 |
| 2.31873 | 0.0118 |
| 2.33493 | 0.0123 |
| 2.35114 | 0.0117 |
| 2.36896 | 0.0121 |
| 2.38516 | 0.0124 |
| 2.40137 | 0.0123 |
| 2.41757 | 0.0131 |
| 2.43539 | 0.0126 |
| 2.4516 | 0.0127 |
| 2.4678 | 0.0124 |
| 2.48563 | 0.0115 |
| 2.50183 | 0.0125 |
| 2.51803 | 0.0118 |
| 2.53423 | 0.0122 |
| 2.55206 | 0.0103 |
| 2.56826 | 0.0103 |
| 2.58447 | 0.01 |
| 2.60229 | 0.01 |
| 2.61849 | 0.0098 |
| 2.6347 | 0.0096 |
| 2.6509 | 0.0102 |
| 2.66872 | 0.0087 |
| 2.68493 | 0.0087 |
| 2.70113 | 0.0091 |
| 2.71896 | 0.0087 |
| 2.73516 | 0.008 |
| 2.75136 | 0.0082 |
| 2.76757 | 0.0091 |
| 2.78539 | 0.0071 |
| 2.80159 | 0.009 |
| 2.8178 | 0.0082 |
| 2.83562 | 0.0085 |
| 2.85182 | 0.0078 |
| 2.86803 | 0.009 |
| 2.88423 | 0.011 |
| 2.90206 | 0.0104 |
| 2.91826 | 0.0101 |
| 2.93446 | 0.0091 |
| 2.95229 | 0.0105 |
| 2.96849 | 0.0115 |
| 2.98469 | 0.0121 |
| 3.0009 | 0.0114 |
| 3.01872 | 0.011 |
| 3.03492 | 0.012 |
| 3.05113 | 0.0111 |
| 3.06895 | 0.0107 |
| 3.08516 | 0.0117 |
| 3.10136 | 0.0125 |
| 3.11756 | 0.0124 |
| 3.13539 | 0.0128 |
| 3.15159 | 0.0127 |
| 3.16779 | 0.0129 |
| 3.18562 | 0.0119 |
| 3.20182 | 0.0124 |
| 3.21802 | 0.0135 |
| 3.23423 | 0.0128 |
| 3.25205 | 0.0118 |
| 3.26826 | 0.0118 |
| 3.28446 | 0.0114 |
| 3.30228 | 0.0104 |
| 3.31849 | 0.0124 |
| 3.33469 | 0.0124 |
| 3.35251 | 0.0122 |

| C18 retention time, min | Borate, pH 8, 37 °C, Overnight |
| --- | --- |
| 0.050231 | 0.0004 |
| 0.0599531 | -0.0003 |
| 0.0761567 | -0.001 |
| 0.0939806 | 0.0012 |
| 0.110184 | -0.0002 |
| 0.126388 | 0.0008 |
| 0.142591 | 0.0002 |
| 0.158795 | 0.0015 |
| 0.174998 | -0.0003 |
| 0.191202 | -0.0001 |
| 0.209026 | 0.0004 |
| 0.225229 | 0.0005 |
| 0.241433 | -0.0001 |
| 0.257636 | -0.0006 |
| 0.27546 | 0.0009 |
| 0.291664 | -0.0007 |
| 0.307867 | -0.0004 |
| 0.325691 | -0.0001 |
| 0.341895 | -0.0001 |
| 0.358098 | -0.0013 |
| 0.374302 | -0.001 |
| 0.392126 | -0.0013 |
| 0.40833 | -0.0011 |
| 0.424533 | -0.0015 |
| 0.442357 | -0.0022 |
| 0.460181 | -0.0014 |
| 0.476384 | -0.002 |
| 0.492588 | -0.0028 |
| 0.510412 | -0.0021 |
| 0.526615 | -0.0023 |
| 0.542819 | -0.0013 |
| 0.559022 | -0.0023 |
| 0.575226 | -0.0035 |
| 0.59143 | -0.0048 |
| 0.607633 | -0.0034 |
| 0.625457 | -0.0033 |
| 0.641661 | -0.0035 |
| 0.657864 | -0.0029 |
| 0.675688 | -0.0041 |
| 0.691892 | -0.0043 |
| 0.708095 | -0.0037 |
| 0.725919 | -0.0032 |
| 0.742123 | -0.0028 |
| 0.759946 | -0.0031 |
| 0.77615 | -0.0032 |
| 0.793974 | -0.0027 |
| 0.810177 | -0.0021 |
| 0.826381 | -0.0027 |
| 0.842585 | -0.0018 |
| 0.858788 | -0.0018 |
| 0.874992 | -0.0004 |
| 0.891195 | -0.0007 |
| 0.909019 | -0.0001 |
| 0.925223 | -0.0004 |
| 0.941426 | -0.0015 |
| 0.95763 | -0.0005 |
| 0.975454 | -0.0007 |
| 0.991657 | -0.0003 |
| 1.00786 | -0.0003 |
| 1.02568 | 0.0002 |
| 1.04189 | 0.0007 |
| 1.05809 | 0.0005 |
| 1.07592 | 0.0003 |
| 1.0905 | 0.0006 |
| 1.1067 | 0.0006 |
| 1.12291 | 0.0009 |
| 1.14073 | -0.0004 |
| 1.15855 | -0.0001 |
| 1.17476 | 0.0001 |
| 1.19096 | 0 |
| 1.20878 | -0.0001 |
| 1.22499 | -0.0004 |
| 1.24119 | 0.0015 |
| 1.25902 | 0.0009 |
| 1.27522 | -0.0004 |
| 1.29142 | -0.0007 |
| 1.30763 | -0.0008 |
| 1.32545 | 0.0002 |
| 1.34165 | 0.0005 |
| 1.35786 | 0.0002 |
| 1.37568 | 0.0013 |
| 1.39189 | -0.0004 |
| 1.40809 | 0.0002 |
| 1.42429 | -0.0003 |
| 1.44212 | -0.0003 |
| 1.45832 | 0.0008 |
| 1.47452 | 0.0009 |
| 1.49235 | 0.0006 |
| 1.50855 | 0 |
| 1.52475 | 0.0004 |
| 1.54096 | 0.0004 |
| 1.55878 | 0.0003 |
| 1.57499 | 0.0012 |
| 1.59119 | 0 |
| 1.60901 | -0.0008 |
| 1.62522 | 0.0001 |
| 1.64142 | 0.0004 |
| 1.65762 | 0.0001 |
| 1.67545 | 0.0003 |
| 1.69165 | 0.0005 |
| 1.70785 | 0.0006 |
| 1.72568 | 0.0009 |
| 1.74188 | -0.001 |
| 1.75808 | 0.0002 |
| 1.77591 | 0.0005 |
| 1.79211 | 0.0004 |
| 1.80832 | -0.0003 |
| 1.82452 | 0.0004 |
| 1.84234 | 0.0009 |
| 1.85855 | 0.0005 |
| 1.87475 | 0.0006 |
| 1.89257 | -0.0007 |
| 1.90878 | 0.0005 |
| 1.92498 | 0.0009 |
| 1.94118 | 0.0018 |
| 1.95901 | 0.0009 |
| 1.97521 | 0.0013 |
| 1.99142 | 0.0017 |
| 2.00924 | 0.0005 |
| 2.02544 | 0.0015 |
| 2.04165 | 0.0008 |
| 2.05785 | 0.0021 |
| 2.07567 | 0.0027 |
| 2.09188 | 0.0023 |
| 2.10808 | 0.0031 |
| 2.12428 | 0.0032 |
| 2.14211 | 0.0032 |
| 2.15831 | 0.0048 |
| 2.17452 | 0.0061 |
| 2.19234 | 0.005 |
| 2.20854 | 0.007 |
| 2.22475 | 0.0081 |
| 2.24257 | 0.0074 |
| 2.25877 | 0.0088 |
| 2.27498 | 0.0086 |
| 2.29118 | 0.0092 |
| 2.30901 | 0.0108 |
| 2.32521 | 0.0107 |
| 2.34141 | 0.0117 |
| 2.35762 | 0.0127 |
| 2.37544 | 0.014 |
| 2.39164 | 0.0142 |
| 2.40785 | 0.0144 |
| 2.42567 | 0.015 |
| 2.44188 | 0.0167 |
| 2.45808 | 0.0171 |
| 2.47428 | 0.0171 |
| 2.49211 | 0.0176 |
| 2.50831 | 0.0182 |
| 2.52451 | 0.0196 |
| 2.54234 | 0.0204 |
| 2.55854 | 0.022 |
| 2.57474 | 0.0229 |
| 2.59257 | 0.0244 |
| 2.60877 | 0.0253 |
| 2.62498 | 0.027 |
| 2.64118 | 0.0281 |
| 2.659 | 0.0298 |
| 2.67521 | 0.0298 |
| 2.69141 | 0.0323 |
| 2.70923 | 0.0324 |
| 2.72544 | 0.0349 |
| 2.74164 | 0.0345 |
| 2.75784 | 0.0359 |
| 2.77567 | 0.0369 |
| 2.79187 | 0.038 |
| 2.80808 | 0.0397 |
| 2.8259 | 0.0402 |
| 2.8421 | 0.0414 |
| 2.85831 | 0.0421 |
| 2.87451 | 0.0422 |
| 2.89233 | 0.0431 |
| 2.90854 | 0.0441 |
| 2.92474 | 0.0437 |
| 2.94257 | 0.0445 |
| 2.95877 | 0.0444 |
| 2.97497 | 0.0446 |
| 2.99118 | 0.0455 |
| 3.009 | 0.0447 |
| 3.0252 | 0.0452 |
| 3.04141 | 0.0471 |
| 3.05761 | 0.0461 |
| 3.07381 | 0.0464 |
| 3.09002 | 0.0471 |
| 3.10784 | 0.0461 |
| 3.12404 | 0.0469 |
| 3.14025 | 0.046 |
| 3.15645 | 0.0467 |
| 3.17265 | 0.0469 |
| 3.19048 | 0.0468 |
| 3.20668 | 0.0464 |
| 3.22289 | 0.0471 |
| 3.24071 | 0.0463 |
| 3.25691 | 0.0465 |
| 3.27312 | 0.0455 |
| 3.28932 | 0.0462 |
| 3.30714 | 0.0463 |
| 3.32335 | 0.0463 |
| 3.33955 | 0.0445 |
| 3.35738 | 0.0455 |

| C18 retention time, min | Borate, pH 8.5, 37 °C, Overnight |
| --- | --- |
| 0.050231 | 0.0018 |
| 0.0631938 | 0.0014 |
| 0.0793974 | 0.0019 |
| 0.0972213 | -0.0005 |
| 0.113425 | 0.0012 |
| 0.129628 | 0.0011 |
| 0.149073 | 0.0009 |
| 0.165276 | -0.0002 |
| 0.18148 | 0.0006 |
| 0.197683 | 0.0001 |
| 0.215507 | 0.0004 |
| 0.231711 | -0.0005 |
| 0.247914 | 0.0007 |
| 0.265738 | -0.0006 |
| 0.281942 | -0.0001 |
| 0.298145 | 0.0004 |
| 0.315969 | -0.0001 |
| 0.332173 | -0.0004 |
| 0.348376 | 0.0003 |
| 0.36458 | 0.0003 |
| 0.382404 | 0.0001 |
| 0.398607 | -0.0007 |
| 0.414811 | 0.0001 |
| 0.432635 | -0.0001 |
| 0.448838 | -0.0015 |
| 0.463422 | -0.0003 |
| 0.479625 | -0.0016 |
| 0.497449 | -0.0008 |
| 0.513653 | -0.0011 |
| 0.529856 | 0.0007 |
| 0.54768 | -0.0008 |
| 0.563883 | -0.0005 |
| 0.580087 | -0.0001 |
| 0.596291 | -0.0008 |
| 0.614115 | -0.0008 |
| 0.630318 | -0.0015 |
| 0.646522 | -0.0012 |
| 0.662725 | -0.0007 |
| 0.680549 | -0.0009 |
| 0.696753 | -0.0015 |
| 0.712956 | -0.0003 |
| 0.73078 | -0.0012 |
| 0.746984 | -0.0008 |
| 0.763187 | -0.0004 |
| 0.779391 | -0.0018 |
| 0.797215 | -0.0017 |
| 0.813418 | -0.0007 |
| 0.829622 | -0.0017 |
| 0.847446 | -0.0007 |
| 0.863649 | -0.0004 |
| 0.879853 | -0.0013 |
| 0.896056 | -0.0018 |
| 0.91388 | -0.0007 |
| 0.930084 | -0.0017 |
| 0.946287 | -0.0012 |
| 0.964111 | -0.0011 |
| 0.980315 | -0.0005 |
| 0.996518 | -0.0015 |
| 1.01434 | -0.0001 |
| 1.03055 | -0.0005 |
| 1.04675 | -0.0005 |
| 1.06295 | -0.0002 |
| 1.08078 | -0.0011 |
| 1.09698 | -0.0012 |
| 1.11318 | -0.0015 |
| 1.12939 | -0.001 |
| 1.14721 | -0.0017 |
| 1.16342 | -0.0018 |
| 1.17962 | -0.002 |
| 1.19744 | -0.001 |
| 1.21365 | 0.0001 |
| 1.22985 | -0.0013 |
| 1.24605 | -0.0026 |
| 1.26388 | -0.0019 |
| 1.28008 | -0.0019 |
| 1.29628 | -0.0021 |
| 1.31411 | -0.0011 |
| 1.33031 | -0.001 |
| 1.34651 | -0.0025 |
| 1.3611 | -0.0021 |
| 1.37892 | -0.0019 |
| 1.39675 | -0.0017 |
| 1.41133 | -0.0022 |
| 1.42915 | -0.0024 |
| 1.44536 | -0.0014 |
| 1.46156 | -0.0015 |
| 1.47776 | -0.0018 |
| 1.49559 | -0.0013 |
| 1.51179 | -0.0017 |
| 1.52799 | -0.0011 |
| 1.54582 | -0.0022 |
| 1.56202 | -0.0009 |
| 1.57823 | -0.0006 |
| 1.59443 | -0.0015 |
| 1.61225 | -0.0004 |
| 1.62846 | -0.0018 |
| 1.64466 | -0.0016 |
| 1.6641 | -0.002 |
| 1.68031 | -0.0014 |
| 1.69651 | -0.0012 |
| 1.71434 | -0.0006 |
| 1.73054 | -0.0004 |
| 1.74674 | -0.0008 |
| 1.76295 | -0.0015 |
| 1.78077 | -0.0018 |
| 1.79697 | -0.0008 |
| 1.81318 | -0.001 |
| 1.831 | -0.0013 |
| 1.8472 | -0.0014 |
| 1.86341 | -0.0001 |
| 1.87961 | -0.0008 |
| 1.89744 | -0.0009 |
| 1.91364 | -0.0012 |
| 1.92984 | -0.0006 |
| 1.94767 | -0.0003 |
| 1.96387 | 0 |
| 1.98007 | 0.0006 |
| 1.99628 | -0.0007 |
| 2.0141 | 0.0005 |
| 2.0303 | 0.0009 |
| 2.04489 | 0.0016 |
| 2.06109 | 0.0019 |
| 2.07891 | 0.0028 |
| 2.09512 | 0.0031 |
| 2.11132 | 0.0042 |
| 2.12915 | 0.006 |
| 2.14535 | 0.006 |
| 2.16155 | 0.0056 |
| 2.17776 | 0.0069 |
| 2.19558 | 0.0089 |
| 2.21178 | 0.0124 |
| 2.22799 | 0.0124 |
| 2.24581 | 0.0131 |
| 2.26202 | 0.0143 |
| 2.27822 | 0.0176 |
| 2.29442 | 0.0191 |
| 2.31225 | 0.0211 |
| 2.32845 | 0.0223 |
| 2.34465 | 0.0239 |
| 2.36248 | 0.0269 |
| 2.37868 | 0.0282 |
| 2.39488 | 0.0293 |
| 2.41109 | 0.0309 |
| 2.42891 | 0.0337 |
| 2.44512 | 0.0357 |
| 2.46132 | 0.0381 |
| 2.47752 | 0.0416 |
| 2.49535 | 0.0442 |
| 2.51155 | 0.0472 |
| 2.52775 | 0.049 |
| 2.54558 | 0.0508 |
| 2.56178 | 0.0531 |
| 2.57798 | 0.0553 |
| 2.59581 | 0.057 |
| 2.61201 | 0.0582 |
| 2.62822 | 0.0624 |
| 2.64442 | 0.0625 |
| 2.66224 | 0.065 |
| 2.67845 | 0.0656 |
| 2.69465 | 0.0688 |
| 2.71247 | 0.0704 |
| 2.72868 | 0.0719 |
| 2.74488 | 0.0724 |
| 2.76109 | 0.0746 |
| 2.77891 | 0.0761 |
| 2.79511 | 0.0783 |
| 2.81132 | 0.0784 |
| 2.82914 | 0.0794 |
| 2.84534 | 0.0812 |
| 2.86155 | 0.0815 |
| 2.87775 | 0.0836 |
| 2.89557 | 0.0833 |
| 2.91178 | 0.0852 |
| 2.92798 | 0.0866 |
| 2.94581 | 0.0855 |
| 2.96201 | 0.0863 |
| 2.97821 | 0.0872 |
| 2.99442 | 0.0865 |
| 3.01224 | 0.0876 |
| 3.02844 | 0.0886 |
| 3.04465 | 0.0884 |
| 3.06247 | 0.0891 |
| 3.07868 | 0.0897 |
| 3.09488 | 0.0897 |
| 3.11108 | 0.0891 |
| 3.12891 | 0.0903 |
| 3.14511 | 0.0902 |
| 3.16131 | 0.0899 |
| 3.17914 | 0.0902 |
| 3.19534 | 0.0902 |
| 3.21154 | 0.0897 |
| 3.22937 | 0.0911 |
| 3.24557 | 0.0914 |
| 3.26178 | 0.0909 |
| 3.27798 | 0.0901 |
| 3.2958 | 0.0912 |
| 3.31201 | 0.0913 |
| 3.32821 | 0.0921 |
| 3.34603 | 0.091 |
| 3.36224 | 0.0913 |

| C18 retention time, min | Borate, pH 9, 37 °C, Overnight |
| --- | --- |
| 0.050231 | -0.0005 |
| 0.0534717 | 0.0003 |
| 0.0696753 | 0.0002 |
| 0.0874992 | 0.0004 |
| 0.103703 | 0.0009 |
| 0.119906 | 0.0018 |
| 0.13611 | 0.0014 |
| 0.153934 | 0.0024 |
| 0.170137 | 0.0021 |
| 0.186341 | 0.0018 |
| 0.202544 | 0.0009 |
| 0.220368 | 0.0023 |
| 0.236572 | 0.0024 |
| 0.252775 | 0.0025 |
| 0.270599 | 0.0027 |
| 0.286803 | 0.0033 |
| 0.303006 | 0.0022 |
| 0.32083 | 0.0034 |
| 0.335413 | 0.0023 |
| 0.351617 | 0.0037 |
| 0.367821 | 0.0027 |
| 0.385645 | 0.0042 |
| 0.401848 | 0.0018 |
| 0.418052 | 0.003 |
| 0.435876 | 0.003 |
| 0.452079 | 0.0016 |
| 0.468283 | 0.0035 |
| 0.484486 | 0.0029 |
| 0.50231 | 0.002 |
| 0.518514 | 0.0033 |
| 0.534717 | 0.0029 |
| 0.552541 | 0.0013 |
| 0.568745 | 0.0025 |
| 0.584948 | 0.0018 |
| 0.601152 | 0.002 |
| 0.618976 | 0.0017 |
| 0.635179 | 0.0031 |
| 0.651383 | 0.0028 |
| 0.669207 | 0.0027 |
| 0.68541 | 0.0024 |
| 0.701614 | 0.0027 |
| 0.717817 | 0.0032 |
| 0.735641 | 0.0021 |
| 0.751845 | 0.0025 |
| 0.768048 | 0.0034 |
| 0.785872 | 0.0034 |
| 0.803696 | 0.0036 |
| 0.8199 | 0.0018 |
| 0.836103 | 0.0028 |
| 0.853927 | 0.0041 |
| 0.870131 | 0.0033 |
| 0.886334 | 0.0027 |
| 0.902538 | 0.0045 |
| 0.920362 | 0.0031 |
| 0.936565 | 0.0027 |
| 0.952769 | 0.0023 |
| 0.970593 | 0.0024 |
| 0.986796 | 0.0029 |
| 1.003 | 0.0021 |
| 1.02082 | 0.0033 |
| 1.03703 | 0.0022 |
| 1.05323 | 0.0034 |
| 1.06943 | 0.0028 |
| 1.08726 | 0.003 |
| 1.10346 | 0.0028 |
| 1.11966 | 0.0015 |
| 1.13587 | 0.0018 |
| 1.15369 | 0.0014 |
| 1.1699 | 0.0015 |
| 1.1861 | 0.0012 |
| 1.20392 | 0.0016 |
| 1.22013 | 0.0018 |
| 1.23633 | 0.0023 |
| 1.25253 | 0.0021 |
| 1.27036 | 0.0017 |
| 1.28656 | 0.0012 |
| 1.30276 | 0.0028 |
| 1.32059 | 0.0023 |
| 1.33679 | 0.0003 |
| 1.353 | 0.002 |
| 1.37082 | 0.002 |
| 1.38702 | 0.0008 |
| 1.40323 | 0.0007 |
| 1.41943 | 0.0018 |
| 1.43725 | 0.0015 |
| 1.45346 | 0.0012 |
| 1.46966 | 0.0022 |
| 1.48749 | 0.0024 |
| 1.50369 | 0.0011 |
| 1.51989 | 0.002 |
| 1.5361 | 0.0024 |
| 1.5523 | 0.0009 |
| 1.5685 | 0.0022 |
| 1.58471 | 0.0011 |
| 1.60091 | 0.0011 |
| 1.61874 | 0.0023 |
| 1.63494 | 0.0023 |
| 1.65114 | 0.0031 |
| 1.66897 | 0.0019 |
| 1.68517 | 0.0016 |
| 1.70137 | 0.0026 |
| 1.71758 | 0.0023 |
| 1.7354 | 0.0031 |
| 1.7516 | 0.0023 |
| 1.76781 | 0.0013 |
| 1.78563 | 0.002 |
| 1.80184 | 0.002 |
| 1.81804 | 0.0021 |
| 1.83424 | 0.0023 |
| 1.85207 | 0.002 |
| 1.86827 | 0.0023 |
| 1.88447 | 0.0023 |
| 1.9023 | 0.0027 |
| 1.9185 | 0.0034 |
| 1.9347 | 0.0026 |
| 1.95091 | 0.0022 |
| 1.96873 | 0.0035 |
| 1.98493 | 0.0026 |
| 2.00114 | 0.0048 |
| 2.01896 | 0.0044 |
| 2.03517 | 0.0059 |
| 2.05137 | 0.0052 |
| 2.06919 | 0.0061 |
| 2.0854 | 0.0071 |
| 2.1016 | 0.0066 |
| 2.1178 | 0.0084 |
| 2.13563 | 0.0092 |
| 2.15183 | 0.0098 |
| 2.16804 | 0.0127 |
| 2.18586 | 0.0132 |
| 2.20206 | 0.0145 |
| 2.21827 | 0.0163 |
| 2.23447 | 0.0186 |
| 2.25229 | 0.0206 |
| 2.2685 | 0.0233 |
| 2.2847 | 0.0241 |
| 2.3009 | 0.0279 |
| 2.31873 | 0.0293 |
| 2.33493 | 0.0302 |
| 2.35114 | 0.0329 |
| 2.36896 | 0.0367 |
| 2.38516 | 0.0382 |
| 2.40137 | 0.0405 |
| 2.41757 | 0.0438 |
| 2.43539 | 0.0448 |
| 2.4516 | 0.0472 |
| 2.4678 | 0.049 |
| 2.48563 | 0.0521 |
| 2.50183 | 0.0548 |
| 2.51803 | 0.0576 |
| 2.53423 | 0.0602 |
| 2.55206 | 0.0621 |
| 2.56826 | 0.0643 |
| 2.58447 | 0.0681 |
| 2.60229 | 0.0699 |
| 2.61849 | 0.0725 |
| 2.6347 | 0.0747 |
| 2.65252 | 0.0752 |
| 2.66872 | 0.0785 |
| 2.68493 | 0.0797 |
| 2.70113 | 0.082 |
| 2.71733 | 0.0829 |
| 2.73354 | 0.0848 |
| 2.74974 | 0.0863 |
| 2.76757 | 0.0887 |
| 2.78377 | 0.09 |
| 2.79997 | 0.0902 |
| 2.81618 | 0.0925 |
| 2.834 | 0.0944 |
| 2.8502 | 0.0951 |
| 2.86641 | 0.0969 |
| 2.88261 | 0.0973 |
| 2.90043 | 0.0984 |
| 2.91664 | 0.0983 |
| 2.93284 | 0.1011 |
| 2.95067 | 0.0998 |
| 2.96687 | 0.1025 |
| 2.98307 | 0.1023 |
| 2.99928 | 0.1034 |
| 3.0171 | 0.1028 |
| 3.0333 | 0.1032 |
| 3.04951 | 0.1037 |
| 3.06733 | 0.1046 |
| 3.08354 | 0.1043 |
| 3.09974 | 0.1058 |
| 3.11594 | 0.1052 |
| 3.13377 | 0.1066 |
| 3.14997 | 0.1064 |
| 3.16617 | 0.1062 |
| 3.184 | 0.1061 |
| 3.2002 | 0.1062 |
| 3.2164 | 0.1061 |
| 3.23261 | 0.1067 |
| 3.25043 | 0.1074 |
| 3.26664 | 0.1078 |
| 3.28284 | 0.1063 |
| 3.30066 | 0.1072 |
| 3.31687 | 0.1061 |
| 3.33307 | 0.1073 |
| 3.34927 | 0.1059 |

| C18 retention time, min | Borate, pH 9.5, 37 °C, Overnight |
| --- | --- |
| 0.050231 | 0.0041 |
| 0.0631938 | 0.0031 |
| 0.0793974 | 0.0017 |
| 0.0972213 | 0.0033 |
| 0.113425 | 0.003 |
| 0.129628 | 0.0035 |
| 0.147452 | 0.0034 |
| 0.163656 | 0.0023 |
| 0.179859 | 0.0032 |
| 0.196063 | 0.0025 |
| 0.213887 | 0.0017 |
| 0.23009 | 0.0034 |
| 0.244674 | 0.0019 |
| 0.262497 | 0.0034 |
| 0.278701 | 0.003 |
| 0.294905 | 0.0015 |
| 0.311108 | 0.0022 |
| 0.328932 | 0.0028 |
| 0.345136 | 0.0026 |
| 0.361339 | 0.0029 |
| 0.379163 | 0.0018 |
| 0.395367 | 0.0026 |
| 0.41157 | 0.0025 |
| 0.427774 | 0.0024 |
| 0.445598 | 0.0023 |
| 0.461801 | 0.0019 |
| 0.478005 | 0.0029 |
| 0.495829 | 0.0034 |
| 0.512032 | 0.0022 |
| 0.528236 | 0.0015 |
| 0.544439 | 0.0027 |
| 0.562263 | 0.0021 |
| 0.578467 | 0.0008 |
| 0.59467 | 0.0015 |
| 0.612494 | 0.0027 |
| 0.628698 | 0.0014 |
| 0.644901 | 0.0015 |
| 0.661105 | 0.0005 |
| 0.678929 | 0.0021 |
| 0.695132 | 0.0008 |
| 0.711336 | 0.0026 |
| 0.72916 | 0.0013 |
| 0.745363 | 0.0018 |
| 0.761567 | 0.0025 |
| 0.77777 | 0.0006 |
| 0.795594 | 0.0007 |
| 0.811798 | 0.0014 |
| 0.828001 | 0.0014 |
| 0.845825 | 0.0032 |
| 0.862029 | 0.0026 |
| 0.878232 | 0.0016 |
| 0.894436 | 0.0016 |
| 0.91226 | 0.0021 |
| 0.928463 | 0.0009 |
| 0.944667 | 0.0022 |
| 0.962491 | 0.0021 |
| 0.978694 | 0.0016 |
| 0.994898 | 0.0023 |
| 1.0111 | 0.0017 |
| 1.02893 | 0.0018 |
| 1.04513 | 0.0011 |
| 1.06295 | 0.0013 |
| 1.08078 | 0.002 |
| 1.09698 | 0.0022 |
| 1.11156 | 0.0011 |
| 1.12939 | 0.0022 |
| 1.14559 | 0.0008 |
| 1.16179 | 0.0011 |
| 1.178 | 0.0012 |
| 1.19582 | 0.001 |
| 1.21203 | 0.0014 |
| 1.22823 | 0.0019 |
| 1.24443 | 0.001 |
| 1.26226 | 0.0015 |
| 1.27684 | 0.0008 |
| 1.29304 | 0.0007 |
| 1.31087 | 0.0017 |
| 1.32707 | 0.0008 |
| 1.34327 | 0.0021 |
| 1.35948 | 0.0016 |
| 1.3773 | 0.0026 |
| 1.3935 | 0.0015 |
| 1.40971 | 0.0018 |
| 1.42753 | 0.0012 |
| 1.44374 | 0.0018 |
| 1.46156 | 0.0013 |
| 1.47776 | 0.002 |
| 1.49559 | 0.0019 |
| 1.51179 | 0.0025 |
| 1.52799 | 0.0024 |
| 1.54582 | 0.0028 |
| 1.56202 | 0.0019 |
| 1.57823 | 0.0024 |
| 1.59443 | 0.0028 |
| 1.61225 | 0.0018 |
| 1.62846 | 0.0021 |
| 1.64466 | 0.003 |
| 1.66248 | 0.0027 |
| 1.67869 | 0.0025 |
| 1.69489 | 0.003 |
| 1.7111 | 0.0017 |
| 1.72892 | 0.0026 |
| 1.74512 | 0.0013 |
| 1.76133 | 0.0026 |
| 1.77915 | 0.0025 |
| 1.79535 | 0.0019 |
| 1.81156 | 0.0023 |
| 1.82776 | 0.0028 |
| 1.84558 | 0.0021 |
| 1.86179 | 0.0024 |
| 1.87799 | 0.002 |
| 1.89582 | 0.0028 |
| 1.91202 | 0.0021 |
| 1.92822 | 0.0034 |
| 1.94605 | 0.0035 |
| 1.96225 | 0.0038 |
| 1.97845 | 0.0031 |
| 1.99466 | 0.004 |
| 2.01248 | 0.0045 |
| 2.02868 | 0.0056 |
| 2.04489 | 0.0066 |
| 2.06109 | 0.0065 |
| 2.07891 | 0.0088 |
| 2.09512 | 0.0101 |
| 2.11132 | 0.0114 |
| 2.12915 | 0.0134 |
| 2.14535 | 0.0144 |
| 2.15993 | 0.0181 |
| 2.17614 | 0.0194 |
| 2.19396 | 0.0223 |
| 2.21016 | 0.0258 |
| 2.22637 | 0.0271 |
| 2.24419 | 0.0313 |
| 2.26039 | 0.0349 |
| 2.2766 | 0.0389 |
| 2.2928 | 0.0442 |
| 2.31063 | 0.0471 |
| 2.32683 | 0.0507 |
| 2.34303 | 0.0554 |
| 2.36086 | 0.0576 |
| 2.37706 | 0.0625 |
| 2.39326 | 0.0637 |
| 2.40947 | 0.0686 |
| 2.42729 | 0.0718 |
| 2.44349 | 0.0738 |
| 2.4597 | 0.0759 |
| 2.47752 | 0.0803 |
| 2.49373 | 0.0813 |
| 2.50993 | 0.084 |
| 2.52613 | 0.0861 |
| 2.54396 | 0.0867 |
| 2.56016 | 0.0888 |
| 2.57637 | 0.0895 |
| 2.59419 | 0.09 |
| 2.61039 | 0.0904 |
| 2.6266 | 0.0901 |
| 2.6428 | 0.0908 |
| 2.66062 | 0.0908 |
| 2.67683 | 0.0903 |
| 2.69303 | 0.0913 |
| 2.71085 | 0.0906 |
| 2.72706 | 0.0898 |
| 2.74326 | 0.091 |
| 2.75947 | 0.0892 |
| 2.77729 | 0.0894 |
| 2.79349 | 0.0891 |
| 2.8097 | 0.0882 |
| 2.82752 | 0.0888 |
| 2.84372 | 0.0872 |
| 2.85993 | 0.086 |
| 2.87613 | 0.0873 |
| 2.89395 | 0.0864 |
| 2.91016 | 0.086 |
| 2.92636 | 0.0866 |
| 2.94419 | 0.0858 |
| 2.96201 | 0.0843 |
| 2.97821 | 0.0848 |
| 2.99442 | 0.0857 |
| 3.01224 | 0.0854 |
| 3.02844 | 0.0855 |
| 3.04465 | 0.085 |
| 3.06247 | 0.0844 |
| 3.07868 | 0.0863 |
| 3.09488 | 0.0855 |
| 3.11108 | 0.0856 |
| 3.12891 | 0.084 |
| 3.14511 | 0.0855 |
| 3.15969 | 0.0858 |
| 3.17752 | 0.0854 |
| 3.19372 | 0.0848 |
| 3.20992 | 0.0863 |
| 3.22613 | 0.0857 |
| 3.24395 | 0.0858 |
| 3.26015 | 0.0863 |
| 3.27636 | 0.0863 |
| 3.29418 | 0.0849 |
| 3.31039 | 0.0862 |
| 3.32659 | 0.0843 |
| 3.34279 | 0.0855 |
| 3.36062 | 0.0857 |

Absorbance (214 nm), AU

| C18 retention time, min | Carbonate, Control |
| --- | --- |
| 0.050231 | -0.0009 |
| 0.0550921 | -0.0013 |
| 0.0712956 | -0.0028 |
| 0.0891195 | -0.0013 |
| 0.105323 | -0.0014 |
| 0.121527 | -0.0022 |
| 0.13773 | -0.0022 |
| 0.155554 | -0.0019 |
| 0.171758 | -0.0014 |
| 0.187961 | -0.0017 |
| 0.205785 | -0.001 |
| 0.221989 | -0.0032 |
| 0.238192 | -0.0007 |
| 0.254396 | -0.0011 |
| 0.27222 | -0.001 |
| 0.288423 | -0.0019 |
| 0.304627 | -0.0019 |
| 0.322451 | -0.0016 |
| 0.338654 | -0.0014 |
| 0.354858 | -0.0028 |
| 0.371061 | -0.0006 |
| 0.388885 | -0.0019 |
| 0.405089 | -0.0022 |
| 0.421292 | -0.0024 |
| 0.439116 | -0.0019 |
| 0.45532 | -0.002 |
| 0.471523 | -0.0026 |
| 0.487727 | -0.0022 |
| 0.505551 | -0.0017 |
| 0.521754 | -0.0012 |
| 0.537958 | -0.0028 |
| 0.555782 | -0.0028 |
| 0.571985 | -0.0028 |
| 0.588189 | -0.002 |
| 0.604392 | -0.002 |
| 0.622216 | -0.0015 |
| 0.63842 | -0.0027 |
| 0.654623 | -0.003 |
| 0.672447 | -0.0016 |
| 0.688651 | -0.0012 |
| 0.704854 | -0.0019 |
| 0.721058 | -0.0023 |
| 0.738882 | -0.0015 |
| 0.755085 | -0.0021 |
| 0.771289 | -0.0021 |
| 0.789113 | -0.0024 |
| 0.805316 | -0.0005 |
| 0.82152 | -0.0013 |
| 0.837724 | -0.001 |
| 0.855547 | -0.0021 |
| 0.871751 | -0.0016 |
| 0.887954 | -0.0009 |
| 0.904158 | -0.0016 |
| 0.920362 | -0.0018 |
| 0.936565 | -0.0024 |
| 0.952769 | -0.0005 |
| 0.970593 | -0.0008 |
| 0.986796 | -0.0013 |
| 1.003 | -0.0012 |
| 1.02082 | -0.0015 |
| 1.03703 | -0.003 |
| 1.05323 | -0.003 |
| 1.06943 | -0.0023 |
| 1.08726 | -0.0016 |
| 1.10346 | -0.0009 |
| 1.11966 | -0.0021 |
| 1.13587 | -0.0019 |
| 1.15369 | -0.002 |
| 1.1699 | -0.0013 |
| 1.1861 | -0.0025 |
| 1.20392 | -0.0017 |
| 1.22013 | -0.0009 |
| 1.23633 | -0.0019 |
| 1.25415 | -0.0014 |
| 1.27036 | 0 |
| 1.28656 | -0.0016 |
| 1.30276 | -0.0015 |
| 1.32059 | -0.0021 |
| 1.33679 | -0.0029 |
| 1.353 | -0.0017 |
| 1.37082 | -0.0004 |
| 1.38702 | -0.0009 |
| 1.40323 | -0.0005 |
| 1.41943 | -0.0004 |
| 1.43725 | -0.0016 |
| 1.45346 | -0.0016 |
| 1.46966 | -0.0025 |
| 1.48749 | -0.0023 |
| 1.50369 | -0.0023 |
| 1.51989 | -0.0014 |
| 1.5361 | -0.0019 |
| 1.55392 | -0.0015 |
| 1.57012 | -0.0016 |
| 1.58633 | -0.0024 |
| 1.60415 | -0.0022 |
| 1.62035 | -0.0018 |
| 1.63656 | -0.0024 |
| 1.65276 | -0.0026 |
| 1.67059 | -0.0017 |
| 1.68679 | -0.0026 |
| 1.70299 | -0.0011 |
| 1.72082 | -0.0021 |
| 1.73702 | -0.0008 |
| 1.75322 | -0.0011 |
| 1.76943 | -0.0033 |
| 1.78725 | -0.0019 |
| 1.80346 | -0.0014 |
| 1.81966 | -0.0019 |
| 1.83748 | -0.0016 |
| 1.85369 | -0.0024 |
| 1.86989 | -0.0011 |
| 1.88609 | -0.0004 |
| 1.90392 | -0.0018 |
| 1.92012 | -0.0012 |
| 1.93632 | 0.0002 |
| 1.95415 | 0.0005 |
| 1.97035 | 0.0007 |
| 1.98493 | -0.0001 |
| 2.00114 | 0.0004 |
| 2.01896 | 0.0011 |
| 2.03517 | 0.0027 |
| 2.05137 | 0.003 |
| 2.06919 | 0.0042 |
| 2.0854 | 0.006 |
| 2.1016 | 0.0076 |
| 2.1178 | 0.0087 |
| 2.13563 | 0.0102 |
| 2.15183 | 0.0119 |
| 2.16804 | 0.0158 |
| 2.18586 | 0.0189 |
| 2.20206 | 0.021 |
| 2.21827 | 0.0246 |
| 2.23447 | 0.0281 |
| 2.25229 | 0.0318 |
| 2.2685 | 0.0364 |
| 2.2847 | 0.0408 |
| 2.3009 | 0.0449 |
| 2.31873 | 0.0484 |
| 2.33493 | 0.0533 |
| 2.35114 | 0.0577 |
| 2.36896 | 0.0613 |
| 2.38516 | 0.0661 |
| 2.40137 | 0.0707 |
| 2.41757 | 0.0733 |
| 2.43539 | 0.0777 |
| 2.4516 | 0.0795 |
| 2.4678 | 0.0823 |
| 2.48563 | 0.0828 |
| 2.50183 | 0.0858 |
| 2.51803 | 0.0862 |
| 2.53423 | 0.0873 |
| 2.55206 | 0.0875 |
| 2.56826 | 0.0863 |
| 2.58447 | 0.0855 |
| 2.60229 | 0.0841 |
| 2.61849 | 0.0842 |
| 2.6347 | 0.0818 |
| 2.6509 | 0.0788 |
| 2.66872 | 0.0767 |
| 2.68493 | 0.0747 |
| 2.70113 | 0.072 |
| 2.71896 | 0.0696 |
| 2.73516 | 0.0666 |
| 2.75136 | 0.0644 |
| 2.76757 | 0.0611 |
| 2.78539 | 0.0584 |
| 2.80159 | 0.0546 |
| 2.8178 | 0.0528 |
| 2.83562 | 0.0512 |
| 2.85182 | 0.0491 |
| 2.86803 | 0.0471 |
| 2.88585 | 0.0457 |
| 2.90206 | 0.0422 |
| 2.91826 | 0.0419 |
| 2.93446 | 0.04 |
| 2.95229 | 0.0386 |
| 2.96849 | 0.0367 |
| 2.98469 | 0.0353 |
| 3.0009 | 0.034 |
| 3.01872 | 0.0339 |
| 3.03492 | 0.0336 |
| 3.05113 | 0.0326 |
| 3.06895 | 0.0309 |
| 3.08516 | 0.0314 |
| 3.10136 | 0.0314 |
| 3.11918 | 0.0303 |
| 3.13539 | 0.0294 |
| 3.15159 | 0.0283 |
| 3.16779 | 0.0286 |
| 3.18562 | 0.029 |
| 3.20344 | 0.0287 |
| 3.21965 | 0.0293 |
| 3.23747 | 0.0292 |
| 3.25367 | 0.0282 |
| 3.26988 | 0.0282 |
| 3.28608 | 0.0279 |
| 3.3039 | 0.0273 |
| 3.32011 | 0.027 |
| 3.33631 | 0.0273 |
| 3.35413 | 0.0272 |

| C18 retention time, min | Carbonate, pH 9.2, 4 °C, 42 h |
| --- | --- |
| 0.050231 | 0.0008 |
| 0.0567124 | 0.0009 |
| 0.072916 | -0.0012 |
| 0.0907399 | -0.0004 |
| 0.106943 | -0.0009 |
| 0.123147 | -0.0008 |
| 0.140971 | -0.0008 |
| 0.157174 | 0 |
| 0.173378 | 0.0003 |
| 0.189581 | 0.0004 |
| 0.207405 | -0.0002 |
| 0.223609 | 0.0005 |
| 0.239813 | -0.0002 |
| 0.256016 | 0.0007 |
| 0.27384 | 0.0002 |
| 0.290044 | 0.0002 |
| 0.306247 | -0.0008 |
| 0.324071 | -0.0001 |
| 0.340274 | -0.0005 |
| 0.356478 | 0.0006 |
| 0.372682 | -0.0008 |
| 0.390506 | 0.0001 |
| 0.406709 | 0.0001 |
| 0.422913 | -0.0007 |
| 0.440737 | 0.0009 |
| 0.45694 | 0.0012 |
| 0.473144 | 0.0011 |
| 0.490968 | -0.0001 |
| 0.507171 | 0.0006 |
| 0.523375 | 0.0006 |
| 0.539578 | 0.0001 |
| 0.559022 | 0 |
| 0.575226 | -0.0003 |
| 0.59143 | 0.0009 |
| 0.607633 | 0 |
| 0.625457 | -0.0004 |
| 0.641661 | 0.0008 |
| 0.657864 | -0.0001 |
| 0.675688 | 0.0004 |
| 0.691892 | -0.0011 |
| 0.708095 | -0.0001 |
| 0.724299 | -0.0005 |
| 0.742123 | 0.0008 |
| 0.758326 | 0.0014 |
| 0.77453 | 0.001 |
| 0.792354 | 0.0002 |
| 0.808557 | 0.0004 |
| 0.824761 | 0.0004 |
| 0.840964 | 0.0002 |
| 0.858788 | 0.0003 |
| 0.874992 | -0.0005 |
| 0.891195 | 0.0006 |
| 0.909019 | 0.0005 |
| 0.925223 | 0.0003 |
| 0.941426 | 0 |
| 0.95763 | -0.0004 |
| 0.975454 | -0.0007 |
| 0.991657 | -0.0004 |
| 1.00786 | 0.0001 |
| 1.02568 | 0.0007 |
| 1.04189 | 0.0002 |
| 1.05647 | 0.0008 |
| 1.07268 | -0.0007 |
| 1.0905 | 0.0004 |
| 1.1067 | -0.0002 |
| 1.12291 | -0.0003 |
| 1.14073 | -0.0005 |
| 1.15693 | 0.0002 |
| 1.17314 | -0.001 |
| 1.18934 | -0.0013 |
| 1.20717 | -0.0005 |
| 1.22337 | 0.0002 |
| 1.23957 | 0.001 |
| 1.2574 | 0.0006 |
| 1.2736 | 0 |
| 1.2898 | -0.0005 |
| 1.30601 | 0 |
| 1.32383 | 0.0008 |
| 1.34003 | -0.0009 |
| 1.35624 | -0.0003 |
| 1.37406 | 0.0002 |
| 1.39027 | 0.0009 |
| 1.40647 | 0.0006 |
| 1.42429 | 0.0013 |
| 1.4405 | -0.0004 |
| 1.4567 | -0.0002 |
| 1.4729 | 0.0003 |
| 1.49073 | 0.0001 |
| 1.50693 | 0.0004 |
| 1.52313 | -0.0005 |
| 1.54096 | 0.0006 |
| 1.55716 | 0.0002 |
| 1.57336 | 0 |
| 1.58957 | -0.0002 |
| 1.60739 | -0.0006 |
| 1.6236 | 0.0005 |
| 1.6398 | 0.0005 |
| 1.656 | 0 |
| 1.67383 | 0.0004 |
| 1.69003 | -0.0003 |
| 1.70623 | 0.001 |
| 1.72406 | -0.0002 |
| 1.74026 | -0.0003 |
| 1.75646 | 0.0007 |
| 1.77267 | 0.0015 |
| 1.79049 | 0.0012 |
| 1.8067 | 0.0008 |
| 1.8229 | 0.0006 |
| 1.84072 | 0.0022 |
| 1.85693 | 0.0012 |
| 1.87313 | 0.0012 |
| 1.88933 | 0.0006 |
| 1.90716 | 0.0003 |
| 1.92336 | -0.0003 |
| 1.93956 | 0.0003 |
| 1.95577 | 0.0011 |
| 1.97197 | 0.0006 |
| 1.98818 | 0.0021 |
| 2.00438 | 0.0029 |
| 2.0222 | 0.0034 |
| 2.03841 | 0.0039 |
| 2.05461 | 0.0042 |
| 2.07243 | 0.0055 |
| 2.08864 | 0.0058 |
| 2.10484 | 0.0081 |
| 2.12105 | 0.0102 |
| 2.13887 | 0.0103 |
| 2.15507 | 0.0144 |
| 2.17128 | 0.0175 |
| 2.1891 | 0.0185 |
| 2.2053 | 0.0222 |
| 2.22151 | 0.0264 |
| 2.23933 | 0.0294 |
| 2.25553 | 0.0342 |
| 2.27174 | 0.0373 |
| 2.28794 | 0.0416 |
| 2.30577 | 0.0438 |
| 2.32197 | 0.0469 |
| 2.33817 | 0.0502 |
| 2.35438 | 0.0561 |
| 2.3722 | 0.0594 |
| 2.3884 | 0.064 |
| 2.40461 | 0.0687 |
| 2.42243 | 0.0726 |
| 2.43863 | 0.0779 |
| 2.45646 | 0.082 |
| 2.47266 | 0.0867 |
| 2.49049 | 0.091 |
| 2.50669 | 0.0971 |
| 2.52289 | 0.101 |
| 2.54072 | 0.106 |
| 2.5553 | 0.1091 |
| 2.5715 | 0.1145 |
| 2.58771 | 0.1194 |
| 2.60553 | 0.1227 |
| 2.62173 | 0.1283 |
| 2.63794 | 0.134 |
| 2.65738 | 0.1367 |
| 2.67359 | 0.1418 |
| 2.68979 | 0.1455 |
| 2.70761 | 0.1498 |
| 2.72382 | 0.1546 |
| 2.74002 | 0.1583 |
| 2.75622 | 0.1624 |
| 2.77405 | 0.1659 |
| 2.79025 | 0.1703 |
| 2.80646 | 0.1731 |
| 2.82428 | 0.1757 |
| 2.84048 | 0.1779 |
| 2.85669 | 0.1796 |
| 2.87289 | 0.1827 |
| 2.89071 | 0.1853 |
| 2.90692 | 0.1875 |
| 2.92312 | 0.1885 |
| 2.94094 | 0.1908 |
| 2.95715 | 0.1931 |
| 2.97335 | 0.195 |
| 2.98956 | 0.1954 |
| 3.00738 | 0.1965 |
| 3.02196 | 0.1984 |
| 3.03817 | 0.1979 |
| 3.05437 | 0.2 |
| 3.07219 | 0.1996 |
| 3.0884 | 0.1999 |
| 3.1046 | 0.2018 |
| 3.12242 | 0.2014 |
| 3.13863 | 0.2011 |
| 3.15483 | 0.2015 |
| 3.17103 | 0.2011 |
| 3.18886 | 0.2031 |
| 3.20506 | 0.2028 |
| 3.22127 | 0.2035 |
| 3.23909 | 0.2021 |
| 3.25529 | 0.2028 |
| 3.2715 | 0.2037 |
| 3.2877 | 0.2019 |
| 3.30552 | 0.2038 |
| 3.32173 | 0.2031 |
| 3.33793 | 0.2036 |
| 3.35575 | 0.2034 |

| C18 retention time, min | Carbonate, pH 9.2, 37 °C, Overnight |
| --- | --- |
| 0.050231 | -0.0046 |
| 0.0648142 | -0.0031 |
| 0.0810177 | -0.0032 |
| 0.0988417 | -0.0045 |
| 0.115045 | -0.0031 |
| 0.131249 | -0.004 |
| 0.149073 | -0.004 |
| 0.163656 | -0.0049 |
| 0.179859 | -0.0032 |
| 0.197683 | -0.0042 |
| 0.213887 | -0.005 |
| 0.23009 | -0.0034 |
| 0.247914 | -0.0031 |
| 0.265738 | -0.0049 |
| 0.281942 | -0.0036 |
| 0.298145 | -0.0028 |
| 0.315969 | -0.0038 |
| 0.332173 | -0.004 |
| 0.348376 | -0.0036 |
| 0.36458 | -0.0041 |
| 0.382404 | -0.0031 |
| 0.398607 | -0.004 |
| 0.414811 | -0.0029 |
| 0.432635 | -0.0034 |
| 0.448838 | -0.0035 |
| 0.463422 | -0.004 |
| 0.479625 | -0.0036 |
| 0.497449 | -0.0026 |
| 0.513653 | -0.0042 |
| 0.529856 | -0.0042 |
| 0.54768 | -0.0044 |
| 0.563883 | -0.0035 |
| 0.580087 | -0.0029 |
| 0.596291 | -0.0037 |
| 0.614115 | -0.0037 |
| 0.630318 | -0.0037 |
| 0.646522 | -0.0036 |
| 0.662725 | -0.0035 |
| 0.680549 | -0.0048 |
| 0.696753 | -0.0042 |
| 0.712956 | -0.0044 |
| 0.73078 | -0.0044 |
| 0.746984 | -0.0059 |
| 0.763187 | -0.0052 |
| 0.779391 | -0.0061 |
| 0.797215 | -0.0041 |
| 0.813418 | -0.0046 |
| 0.829622 | -0.0051 |
| 0.847446 | -0.0039 |
| 0.863649 | -0.0049 |
| 0.879853 | -0.0057 |
| 0.896056 | -0.0051 |
| 0.91388 | -0.0055 |
| 0.930084 | -0.0047 |
| 0.946287 | -0.0053 |
| 0.964111 | -0.0054 |
| 0.980315 | -0.0055 |
| 0.996518 | -0.0056 |
| 1.0111 | -0.0051 |
| 1.02893 | -0.0057 |
| 1.04513 | -0.0064 |
| 1.06295 | -0.006 |
| 1.08078 | -0.0053 |
| 1.09698 | -0.0054 |
| 1.11318 | -0.0067 |
| 1.13101 | -0.0051 |
| 1.14721 | -0.0052 |
| 1.16342 | -0.0055 |
| 1.17962 | -0.0048 |
| 1.19744 | -0.0056 |
| 1.21365 | -0.0064 |
| 1.22985 | -0.007 |
| 1.24767 | -0.0065 |
| 1.26388 | -0.006 |
| 1.28008 | -0.0062 |
| 1.29628 | -0.0061 |
| 1.31411 | -0.0054 |
| 1.33031 | -0.0053 |
| 1.34651 | -0.006 |
| 1.36272 | -0.0055 |
| 1.38054 | -0.0056 |
| 1.39675 | -0.0055 |
| 1.41295 | -0.006 |
| 1.43077 | -0.0061 |
| 1.44698 | -0.0062 |
| 1.46318 | -0.006 |
| 1.47938 | -0.0069 |
| 1.49721 | -0.0051 |
| 1.51341 | -0.0062 |
| 1.52961 | -0.0058 |
| 1.54744 | -0.0052 |
| 1.56364 | -0.0047 |
| 1.57985 | -0.0047 |
| 1.59605 | -0.006 |
| 1.61387 | -0.0058 |
| 1.63008 | -0.0039 |
| 1.64628 | -0.0057 |
| 1.6641 | -0.0064 |
| 1.68031 | -0.0054 |
| 1.69651 | -0.0048 |
| 1.71271 | -0.0045 |
| 1.73054 | -0.0057 |
| 1.74674 | -0.0052 |
| 1.76295 | -0.0053 |
| 1.78077 | -0.006 |
| 1.79697 | -0.005 |
| 1.81156 | -0.0062 |
| 1.82776 | -0.0064 |
| 1.84558 | -0.0067 |
| 1.86179 | -0.0064 |
| 1.87799 | -0.0061 |
| 1.89582 | -0.0059 |
| 1.91202 | -0.0051 |
| 1.92822 | -0.0052 |
| 1.94443 | -0.0053 |
| 1.96225 | -0.0052 |
| 1.97845 | -0.0046 |
| 1.99466 | -0.0044 |
| 2.01248 | -0.0052 |
| 2.02868 | -0.0027 |
| 2.04489 | -0.0027 |
| 2.06109 | -0.0014 |
| 2.07891 | 0.0003 |
| 2.09512 | 0.0002 |
| 2.11132 | 0.003 |
| 2.12915 | 0.0059 |
| 2.14535 | 0.0081 |
| 2.16155 | 0.0088 |
| 2.17776 | 0.0124 |
| 2.19558 | 0.0147 |
| 2.21178 | 0.0176 |
| 2.22799 | 0.0208 |
| 2.24581 | 0.0239 |
| 2.26202 | 0.0286 |
| 2.27822 | 0.0308 |
| 2.29442 | 0.0368 |
| 2.31225 | 0.0399 |
| 2.32845 | 0.0451 |
| 2.34465 | 0.0478 |
| 2.36248 | 0.0522 |
| 2.37868 | 0.0551 |
| 2.39488 | 0.06 |
| 2.41109 | 0.0624 |
| 2.42891 | 0.0629 |
| 2.44512 | 0.0661 |
| 2.46132 | 0.0674 |
| 2.47914 | 0.0702 |
| 2.49535 | 0.0719 |
| 2.51155 | 0.0716 |
| 2.52775 | 0.0718 |
| 2.54558 | 0.0716 |
| 2.56178 | 0.0711 |
| 2.57798 | 0.0709 |
| 2.59581 | 0.0687 |
| 2.61201 | 0.0681 |
| 2.62822 | 0.0662 |
| 2.64442 | 0.0645 |
| 2.66224 | 0.0629 |
| 2.67845 | 0.0611 |
| 2.69465 | 0.0595 |
| 2.71247 | 0.0578 |
| 2.72868 | 0.0564 |
| 2.74488 | 0.0552 |
| 2.76109 | 0.0529 |
| 2.77891 | 0.0512 |
| 2.79511 | 0.0486 |
| 2.81132 | 0.0471 |
| 2.82914 | 0.0455 |
| 2.84534 | 0.0446 |
| 2.86155 | 0.044 |
| 2.87775 | 0.0425 |
| 2.89557 | 0.0412 |
| 2.91178 | 0.0393 |
| 2.92798 | 0.0387 |
| 2.94581 | 0.0392 |
| 2.96201 | 0.0375 |
| 2.97821 | 0.036 |
| 2.99442 | 0.0381 |
| 3.01224 | 0.0357 |
| 3.02844 | 0.0367 |
| 3.04465 | 0.0366 |
| 3.06247 | 0.0342 |
| 3.07868 | 0.0339 |
| 3.09488 | 0.0345 |
| 3.11108 | 0.035 |
| 3.12891 | 0.0343 |
| 3.14511 | 0.0343 |
| 3.16131 | 0.0347 |
| 3.17914 | 0.0342 |
| 3.19534 | 0.0334 |
| 3.21154 | 0.0333 |
| 3.22775 | 0.0348 |
| 3.24557 | 0.0348 |
| 3.26015 | 0.0336 |
| 3.27636 | 0.034 |
| 3.29418 | 0.0341 |
| 3.31039 | 0.0334 |
| 3.32659 | 0.0347 |
| 3.34279 | 0.0331 |
| 3.36062 | 0.0341 |

**Figure S7. Full chromatograms of the samples in Figure 12**. Both the peptide peaks and the large carrier protein peaks following them in the gradient are shown. Although all samples had the same initial protein concentration, samples other than borate pH 8.5 and pH 9 showed some reduction in the amount of the cleaved carrier, potentially due to aggregation of the uncleaved protein precursor.

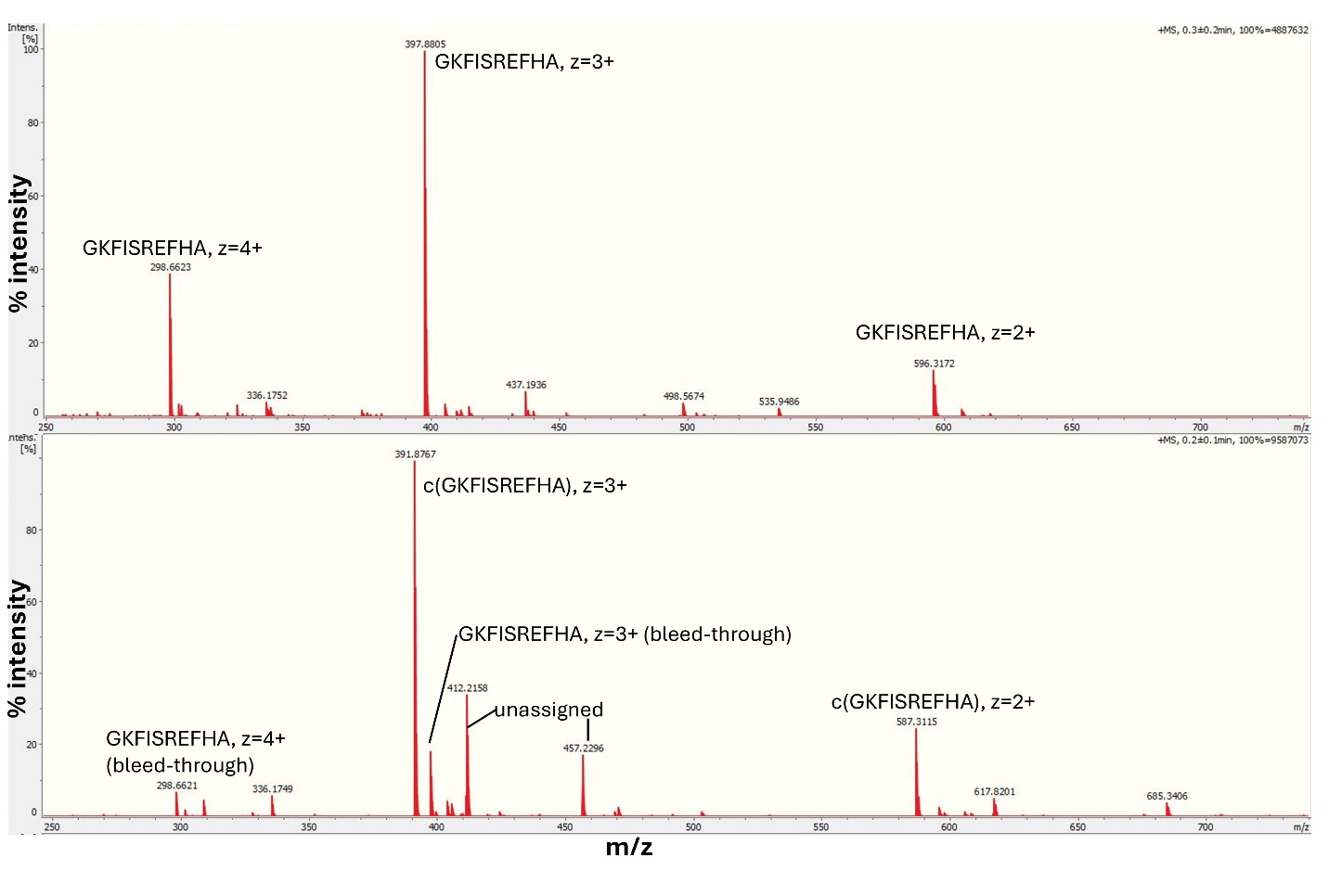

**Figure S8. Direct-injection mass spectrometry of the linear and cyclic product peaks from Figure 12b.** The top panel shows the mass spectrum of the “linear” HPLC peak, and the bottom panel shows the mass spectrum of the “cyclic” HPLC peak. The peaks were isotopically resolved on the Burker Impact II instrument, and we manually confirmed their charge states. For the linear peak, these were: z = +4 for the 298.66 m/z peak; z = +3 for the 397.88 m/z peak; z = +2 for the 596.32 m/z peak. For the cyclic product, these were: z = +3 for the 391.88 m/z peak and z = +2 for the 587.31 m/z peak. c(GKFISREFHA) indicates the cyclized form of the peptide. The most prominent unassigned peaks were 412.22 (z = +3) and 457.23 (z = +3).
